# MEKK3-TGFβ crosstalk regulates inward arterial remodeling

**DOI:** 10.1101/2021.08.19.456893

**Authors:** Hanqiang Deng, Yanying Xu, Xiaoyue Hu, Zhen W. Zhuang, Yuzhou Chang, Yewei Wang, Aglaia Ntokou, Martin A. Schwartz, Bing Su, Michael Simons

**Affiliations:** Yale Cardiovascular Research Center, Department of Internal Medicine, Yale University School of Medicine, New Haven, Connecticut, USA; Shanghai Institute of Immunology, Department of Microbiology and Immunology, and the Ministry of Education Key Laboratory of Cell Death and Differentiation, Shanghai Jiao Tong University School of Medicine, Shanghai, China; Department of Environmental Health Sciences, Yale School of Public Health, New Haven, Connecticut, USA; Department of Cell Biology, Yale University School of Medicine, New Haven, Connecticut, USA; Department of Biomedical Engineering, Yale University, New Haven, Connecticut, USA

**Author notes:** Co-first authors (Hanqiang Deng, Yanying Xu). Co-senior authors (Bing Su, Michael Simons). Address correspondence to, Prof. Michael Simons, Yale University School of Medicine 300 George St, Suite 773, New Haven, CT 06511.

**Keywords:** pulmonary arterial hypertension, systemic hypertension, atherosclerosis, inward arterial remodeling

## Abstract

Arterial remodeling is an important adaptive mechanism that maintains normal fluid shear stress in a variety of physiologic and pathologic conditions. Inward remodeling, a process that leads to reduction in arterial diameter, plays a critical role in progression of such common diseases as hypertension and atherosclerosis. Yet despite its pathogenic importance, molecular mechanisms controlling inward remodeling remain undefined. Mitogen-activated protein kinases (MAPKs) perform a number of functions ranging from control of proliferation to migration and cell fate transitions. While the MAPK ERK1/2 signaling pathway has been extensively examined in the endothelium, less is known about the role of the MEKK3/ERK5 pathway in vascular remodeling. To better define the role played by this signaling cascade, we studied the effect of endothelial-specific deletion of its key upstream MAP3K, MEKK3, in adult mice. The gene’s deletion resulted in a gradual inward remodeling of both pulmonary and systematic arteries leading to spontaneous hypertension in both vascular circuits and accelerated progression of atherosclerosis in hyperlipidemic mice. Molecular analysis revealed activation of TGFβ signaling both in vitro and in vivo. Endothelial-specific TGFβR1 knockout prevented inward arterial remodeling in MEKK3 endothelial knockout mice. These data point to the unexpected participation of endothelial MEKK3 in regulation of TGFβR1-Smad2/3 signaling and inward arterial remodeling in artery diseases.

**Significance:** Inward remodeling of arteries to reduce lumen diameter is a major factor in disease progression and morbidity in multiple vascular diseases, including hypertension and atherosclerosis. However, molecular mechanisms controlling inward arterial remodeling remain largely undefined. In this study, we identify endothelial MEKK3 as an unexpected regulator of inward remodeling via inhibition of TGFβ-Smad2/3 signaling. Genetic deletion of MEKK3 in adult endothelium results in induction of TGFβ-Smad2/3 signaling, endothelial-to-mesenchymal transition and inward remodeling in both pulmonary and arterial circuits. The latter process results in pulmonary and systemic hypertension and accelerates atherosclerosis. These results provide a new basis for understanding the inward artery remodeling that leads to reduced blood flow to affected tissues and exacerbates hypertension in vascular disease.

## Introduction

Vascular remodeling is an adaptive process that matches arterial diameter to blood flow requirements that serves an important role in both physiologic and pathological processes (1, 2). While outward remodeling, induced by increased flow, is well understood, little is known about molecular mechanisms underlying inward remodeling, a process that leads to a reduction in arterial diameter (3, 4). Mitogen activated protein kinases (MAPKs) are central to cells interactions with their environment with many chemical, physical and biological cues processed via this family of enzymes (5). Their specificities, which are tightly associated with their physiological outcomes, are determined mainly through a family of upstream MAPK kinase kinases (MAP3Ks). One member of this family, MEKK3, has been receiving prominent recent attention, in part because of its key role in vascular development (6) and in pathogenesis of cerebral cavernous malformations (CCMs). The kinase’s importance is underlined by a fact that both a decrease and increase in its expression (or activity) lead to profound abnormalities. Thus, increased MEKK3 activity, due to impaired inhibitory interaction with CCM2, leads to CCM formation, in part due to activation of KLF2/KLF4 transcription (7, 8). It also results in abnormal development of other organs, including the heart (9). On the other hand, loss of MEKK3 expression during embryonic or early postnatal development is associated gross abnormalities in vascular development (6).

MEKK3 functions by regulating a signaling cascade leading to activation of its principal targets ERK5 but also of ERK1/2. Endothelial-specific knockout of either MEKK3 itself (10) or its target ERK5 is embryonically lethal (11) as is inactivation of both ERK1 and ERK2 genes (12). Importantly, MEK5/ERK5 pathway is activated, via MEKK3, by laminar fluid shear stress (FSS) (13, 14) and by statins (15), thus suggesting that this signaling cascade is involved in maintenance of normal vascular homeostasis. Indeed, ERK5 activation has been reported to induce a protective phenotype in endothelial cells (ECs) that was associated, in part, with a reduced ECs migration (16) and reduced apoptosis (17) while ERK5 inactivation interferes with anti-inflammatory and anti-apoptotic effects of laminar shear stress (18). However, what endothelial MEKK3 role is in the adult vasculature has not been established. To answer this question, we used a genetic strategy to selectively inactivate MEKK3 in ECs in adult mice and then examined the effects of this deletion.

We find that when induced in adult mice, endothelial-specific deletion of MEKK3 results in a progressive inward remodeling of both pulmonary and systematic vasculatures leading to primary pulmonary and systemic arterial hypertension. Mechanistically, this was linked to activation of endothelial TGFβ signaling and both phenotypes were rescued by an endothelial-specific TGFβR1 knockout. These findings point to a novel molecular pathway controlling inward remodeling and to a critical role MEKK3 plays in regulation of TGFβ signaling.

## Results

### Endothelial MEKK3 knockout induces pulmonary and systemic hypertension

To induce endothelial-specific deletion of MEKK3, previously reported mice with a floxed *Mekk3* gene (19) were crossed with *Cdh5-CreER*^*T2*^ mouse line as described early (10). At 6 weeks of age, mice homozygous for the floxed allele and carrying the *Cdh5-CreER*^*T2*^ transgene (*Cdh5-CreER^*T2*^ Mekk3^fl^*^/*fl*^, hereafter referred to as *Mekk3*^iECKO^) were treated with tamoxifen to induce *mekk3* deletion. *Cdh5-CreER*^*T2*^ negative littermates served as controls. Quantitative polymerase chain reaction (qPCR) analysis of primary lung ECs isolated from *Mekk3*^iECKO^ mice showed 88% decrease of MEKK3 RNA level, but no difference in aortic smooth muscle cells and cardiomyocytes compared to controls (Fig. S1A).

Unlike the previously reported embryonic and neonatal *Mekk3* deletion that led to rapid mortality (6, 10), no deaths were observed in the adult mice up to 5 months post *Mekk3* deletion. However, when sacrificed one month after deletion induction, *Mekk3*^iECKO^ mice displayed a prominent right (RV) and left ventricular (LV) hypertrophy (Fig. 1A and Fig. S1B). To gain a better appreciation for this phenotype, we carried out a time course analysis of changes in RV and LV wall mass. There was a progressive increase in the weight of both ventricles after *Mekk3* deletion induction (Fig. 1 B-D). To determine the cause of this hypertrophy, we measured RV, LV and systolic blood pressures using 1.4F Millar catheters. In agreement with the morphometric data, there was a progressive increase in both RV and LV as well as systemic blood pressure (Fig. 1 E-I and Fig. S1 C-E), pointing to the hemodynamic origin of myocardial hypertrophy in these mice. We also checked heart rate and electrocardiogram (ECG), showing no significant difference between controls and *Mekk3*^iECKO^ mice (Fig. S2 A and B). Since MEKK2 and MEKK3 are highly homologous and both of them can regulate ERK5 activity, we also checked the pressure in previously reported *Mekk2*^−/−^ mice (20). At six months old, western blotting analysis of the lung lysates confirmed the deletion of MEKK2 (Fig. S3A). RV and LV systolic pressure showed no significant difference between controls and *Mekk2*^−/−^ mice (Fig. S3 B and C).

**Figure 1.**
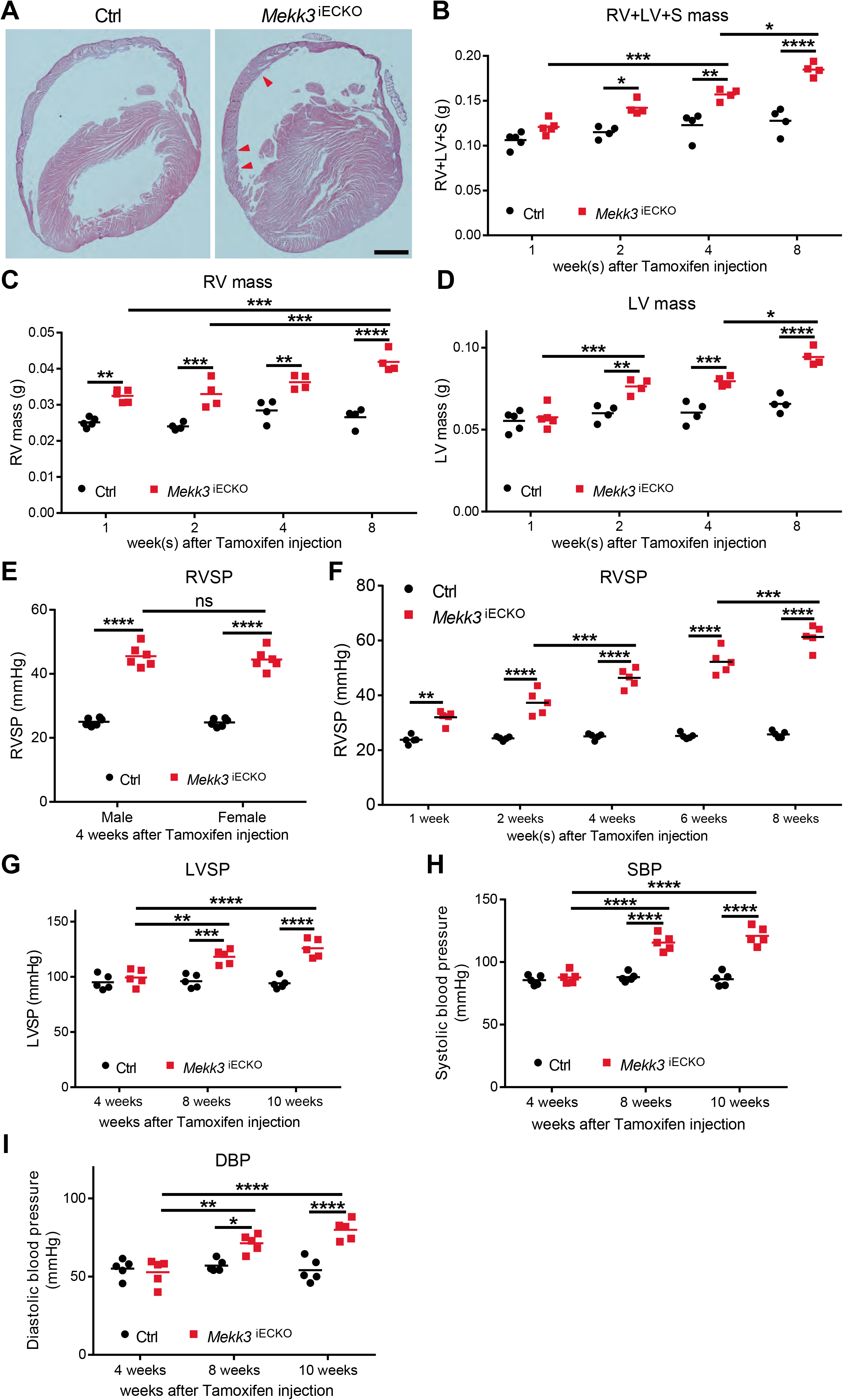
Deletion of endothelial MEKK3 in adult mice induces pulmonary and systemic hypertension. (**A**) Representative H&E staining of Ctrl and *Mekk3*^iECKO^ mice hearts at 4 weeks after tamoxifen injection. Arrowheads point to the right ventricle hypertrophy. Scale bar: 1000μm. (**B-D**) Time course analysis of Right ventricle (RV), left ventricle (LV) and septum (S) mass for Ctrl and *Mekk3*^iECKO^ mice at indicated time points, n=5 at 1 week; n=4 at 2, 4 and 8 weeks, all mice are male. (**E**) Right ventricular systolic pressure (RVSP) of male and female Ctrl and *Mekk3*^iECKO^ mice at 4 weeks after tamoxifen injection, showing no difference between male and female. N=6 mice for per genotype. (**F**) Time course analysis of RVSP for Ctrl and *Mekk3*^iECKO^ mice at indicated time points, n=5 mice (3 male and 2 female) for each time point. (**G-I**) Time course analysis of Left ventricle systolic pressure (LVSP), Systolic blood pressure (SBP) and Diastolic blood pressure (DBP) from Ctrl and *Mekk3*^iECKO^ mice at indicated time points, n=5 male mice for each time point. Bars represent the mean. \**P* < 0.05, \*\**P* < 0.01, \*\*\**P* < 0.001, \*\*\*\**P* < 0.0001, ns: not significant, calculated by two-way ANOVA with Tukey’s multiple comparison tests.

### Endothelial MEKK3 knockout induces inward arterial remodeling

To discern the reason for increased pulmonary and systemic blood pressures in *Mekk3*^iECKO^ mice, we first analyzed the pulmonary vasculature. Micro-CT (mCT) of the lungs showed a large increase in arterial vessel density that was particularly prominent in the periphery (Fig. 2A). Immunohistochemical analysis of whole lung sections stained with a smooth muscle actin (SMA) antibody confirmed this finding, demonstrating a dramatic increase in arterial vascularization that extended throughout the entire lung field (Fig. 2B). Examination of distal pulmonary fields showed a marked increase in smooth muscle coverage of distal pulmonary arterial vessels (Fig. 2C).

**Figure 2.**
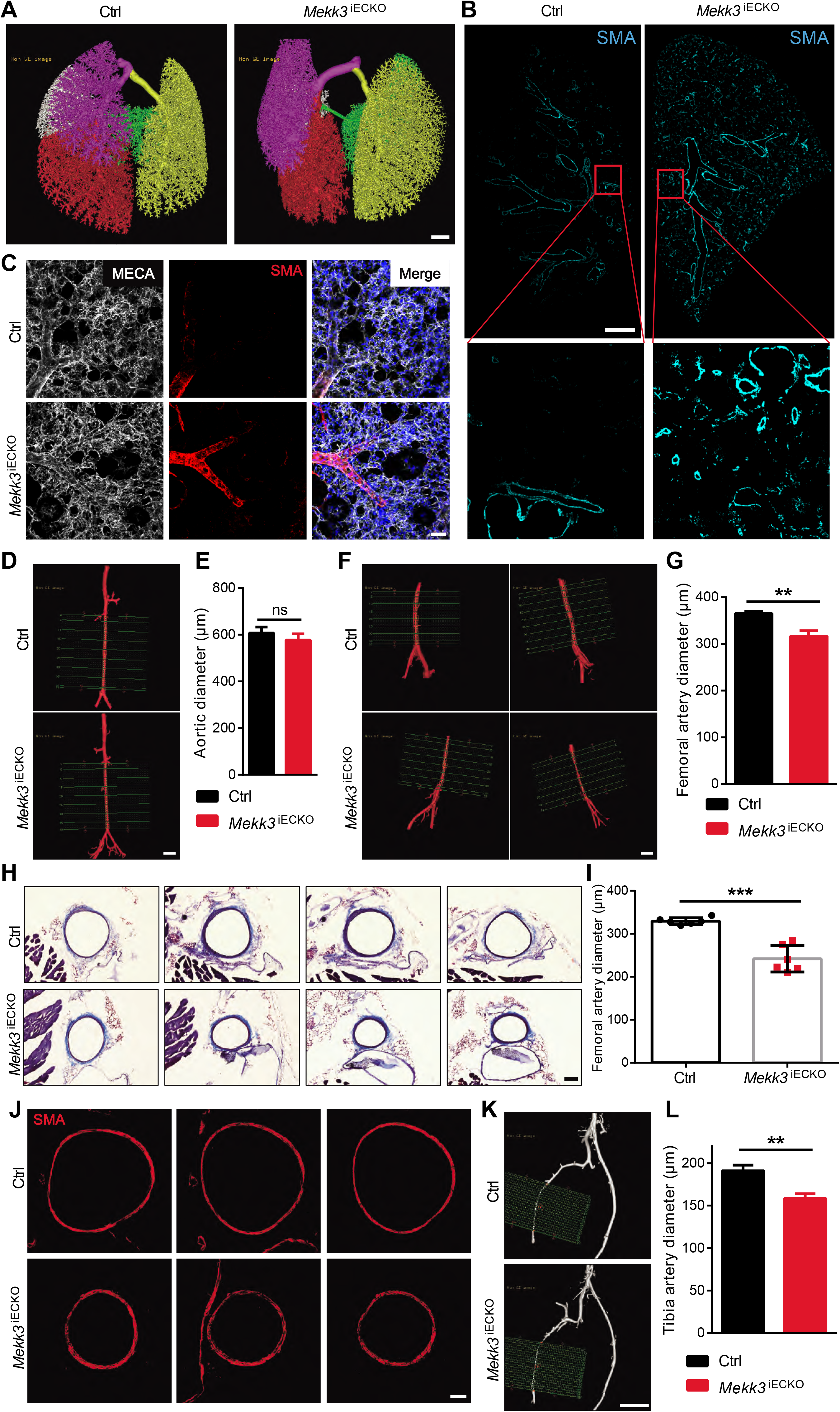
Endothelial MEKK3 deletion leads to inward vascular remolding. (**A**) Representative micro-CT images of Ctrl and *Mekk3*^iECKO^ entire lungs at 6 weeks after tamoxifen injection. Scale bar: 1000μm. (**B**) Representative SMA immunostaining in Ctrl and *Mekk3*^iECKO^ entire lung sections. Scale bar: 1mm. (**C**) Representative higher magnification images of SMA staining of Ctrl and *Mekk3*^iECKO^ lungs. Scale bar: 50μm. (**D**) Representative micro-CT images of Ctrl and *Mekk3*^iECKO^ aorta. Scale bar: 1000μm. (**E**) Quantification of aortic diameter (n=3 male mice). (**F**) Representative micro-CT images of Ctrl and *Mekk3*^iECKO^ femoral artery. Scale bar: 1000μm. (**G**) Quantification of femoral artery diameter for Ctrl (n=6 male mice) and *Mekk3*^iECKO^ mice (n=5 male mice) at 8 weeks after tamoxifen injection. (**H**) Masson Trichrome staining of Ctrl and *Mekk3*^iECKO^ femoral artery sections. Scale bar: 100μm. (**I**) Quantification of femoral artery diameter, n=6 male mice per group. Data represent mean ± SD. (**J**) SMA immunostaining in Ctrl and *Mekk3*^iECKO^ femoral artery sections. Scale bar: 50μm. (**K**) Representative micro-CT images of Ctrl and *Mekk3*^iECKO^ tibia artery. Scale bar: 1000μm. (**L**) Quantification of tibia artery diameter for Ctrl (n=6 male mice) and *Mekk3*^iECKO^ mice (n=5 male mice) at 8 weeks after tamoxifen injection. ns: not significant. \*\**P* < 0.01, \*\*\**P* < 0.001, calculated by unpaired *t*-test.

Since the *Mekk3*^iECKO^ mice also develop systemic hypertension, we next examined peripheral arterial vasculature. While there were no significant changes in mCT-determined aorta diameters between control and *Mekk3*^iECKO^ mice (Fig. 2 D and E), diameters of femoral (Fig. 2 F and G) and tibial (Fig. 2 K and L) arteries were significantly reduced in *Mekk3*^iECKO^ mice. To further confirm these findings, we carried out morphometric analysis of femoral artery sections. In agreement with mCT data, quantification of trichrome-stained sections showed a significant reduction of femoral artery diameter in *Mekk3*^iECKO^ compared to control mice (Fig. 2 H and I). Moreover, SMA staining of femoral artery sections also showed reduction of femoral artery diameter in *Mekk3*^iECKO^ compared to control mice (Fig. 2J).

### Loss of MEKK3 in endothelial cells activates TGFβR signaling and induces EndMT

The observed changes in pulmonary and systemic vasculatures in *Mekk3*^iECKO^ mice point to inward remodeling in medium and smaller size arteries. To obtain molecular insight into this process, we carried out bulk RNA-seq analysis of endothelial cell gene expression at 4 days after MEKK3 knock-down in human umbilical vein endothelial cells (HUVECs). While expression of some of the endothelial genes such as endothelial nitric oxide synthase (eNOS) declined, and some (VEGFR2, VE-cadherin) remained unchanged, there was a pronounced increase in expression of genes associated with endothelial-to-mesenchymal transition (EndMT) including TGFβ receptors 1 and 2 (TGFβR1 and 2), TGFβ2 ligand, SM22α and SMA and extracellular matrix encoding genes fibronectin (FN1) and collagen-1 (COL3A1) as well as a large number of pro-inflammatory genes including matrix metalloproteinases (MMP), chemokines and chemokine receptors, and interleukins (Fig. 3A). These changes in expression were confirmed by qPCR analysis (Fig. 3B), Western blotting (Fig. 3 C and D), immunocytochemistry (Fig. 3 E and F and Fig. S4A) and ELISA (Fig. 3G). MEKK3 knockdown in human pulmonary artery endothelial cells (HPAECs) produced similar results (Fig. S5 A and B).

**Figure 3.**
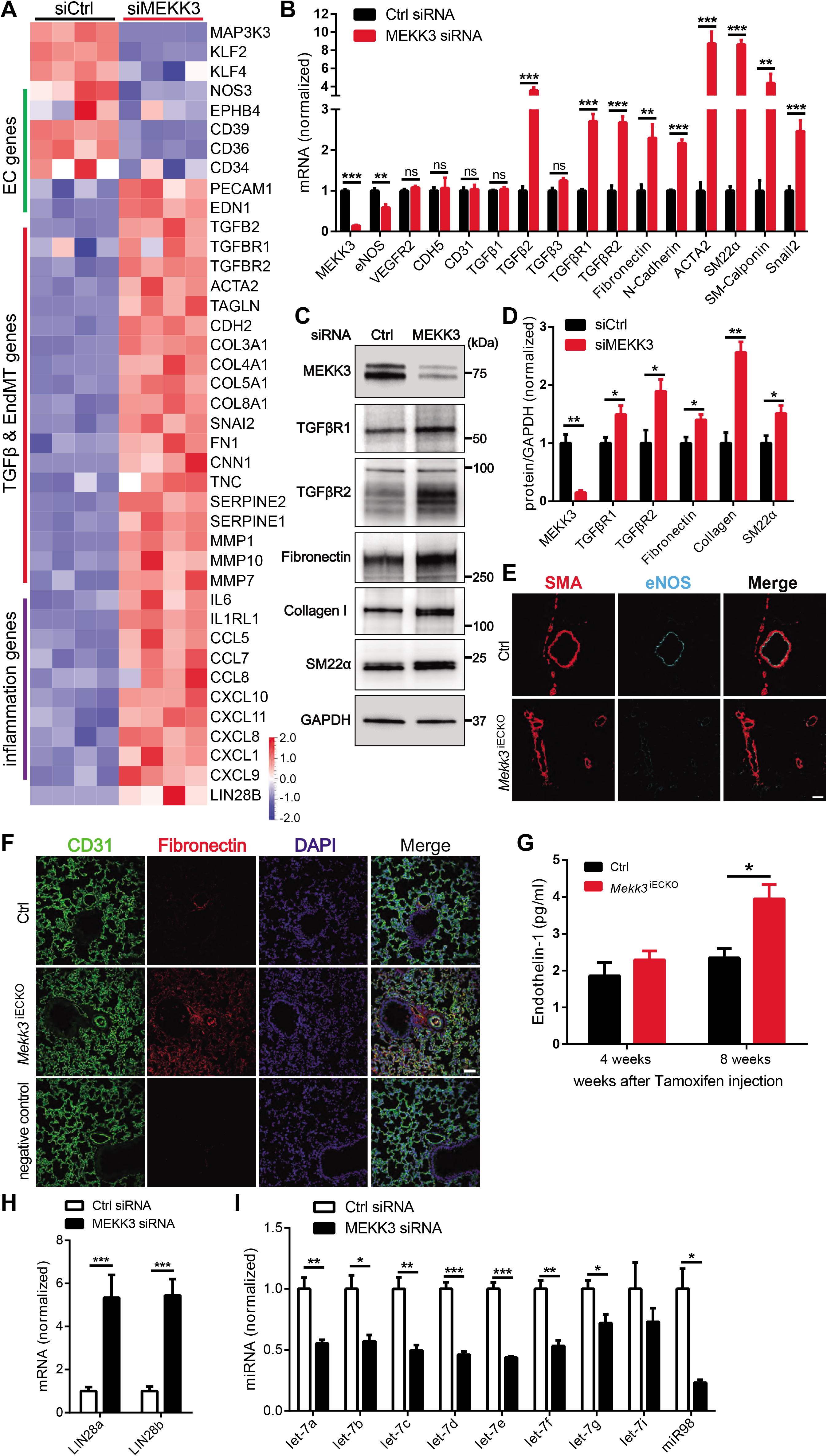
Loss of MEKK3 in ECs induces TGFβ signaling. (**A**) Bulk RNA-seq analysis of Human Umbilical Vein Endothelial Cells (HUVECs) treated with Ctrl or MEKK3 siRNA. Heatmap shows EC, TGFβ, EndMT genes and inflammatory genes. N=4 samples for each group. (**B**) Q-PCR analysis of TGFβ pathway and EndMT markers expression in HUVECs treated with Ctrl or MEKK3 siRNA (n=3). Data represent mean ± SD. (**C-D**) Representative western blot and densitometric quantification of MEKK3 (n=3), TGFβR1 (n=4), TGFβR2 (n=3), fibronectin (n=3), collagen (n=3) and SM22α (n=3) expression in HUVECs treated with Ctrl or MEKK3 siRNA. Data represent mean ± SEM. (**E**) Immunostaining of eNOS and SMA in Ctrl and *Mekk3*^iECKO^ lungs. Scale bar: 20μm. (**F**) Immunostaining of Fibronectin and CD31 in Ctrl and *Mekk3*^iECKO^ lungs. Scale bar: 50μm. (**G**) Circulating endothelin-1 concentration in plasma of Ctrl and *Mekk3*^iECKO^ mice at 4 and 8 weeks after tamoxifen injection (n=4 male mice per group). (**H**) Q-PCR analysis of LIN28a and LIN28b (n=4) expression in HUVECs treated with Ctrl or MEKK3 siRNA. Data represent mean ± SD. (**I**) Q-PCR analysis of let-7 miRNA family (n=3) expression in HUVECs treated with Ctrl or MEKK3 siRNA. ns: not significant. \**P* < 0.05, \*\**P* < 0.01, \*\*\**P* < 0.001, calculated by unpaired *t*-test (B, D, H and I), and two-way ANOVA with Sidak’s multiple comparison tests (G).

Since EndMT induction has been linked to changes in let-7 and Lin28 expression, we used qPCR to quantify expression of these gene families. In agreement with the expression data, we observed an increase in Lin28a and Lin28b level and a corresponding decrease in all let-7 family members’ expression (Fig. 3 H and I). Furthermore, in agreement with the in vitro data, immunocytochemical analysis of Lin28 expression in the lung tissue showed a large increase in its expression in the pulmonary vasculature (Fig. S5C).

Recent investigations have suggested that ERK1/2 activation, especially by FGFs, is central to maintenance of cellular let-7 levels (21–23). Therefore, we tested the effect of MEKK3 knockdown in HUVECs in vitro on FGF2 and VEGF-A165 induced ERK1/2 activation. MEKK3 knockdown reduced the ability of both growth factors to induce ERK1/2 phosphorylation. However, it was far more prominent for FGF2 compared to VEGF-A165 (Fig. S6 A and B). Finally, we tested if MEKK3 is involved in BMP signaling. There was no significant alteration in BMP6 or BMP9-induced Smad-1/5/9 phosphorylation. (Fig. S6 C and D). Also, MEKK3 knockdown did not affect FGFR1 expression (Fig. S6 E and F).

Since we observed a decreased diameter of femoral arteries in *Mekk3*^iECKO^ mice, we next examined the effect of this narrowing on blood flow recovery after ligation of the common femoral artery (hindlimb ischemia model). *Mekk3*^iECKO^ mice exhibited a significantly reduced blood flow recovery compared to controls (Fig. 4 A and B). In agreement with the flow data, we also observed a decreased eNOS expression (Fig. 4 C) in parallel with increased expression of TGFβ and p-Smad3 (Fig. 4 D) in *Mekk3*^iECKO^ gracilis muscle blood vessels. Lastly, we observed a highly significant decrease in the diameter of arterial blood vessels in *Mekk3*^iECKO^ compared to controls (Fig. 4 E), indicating inward arterial remodeling.

**Figure 4.**
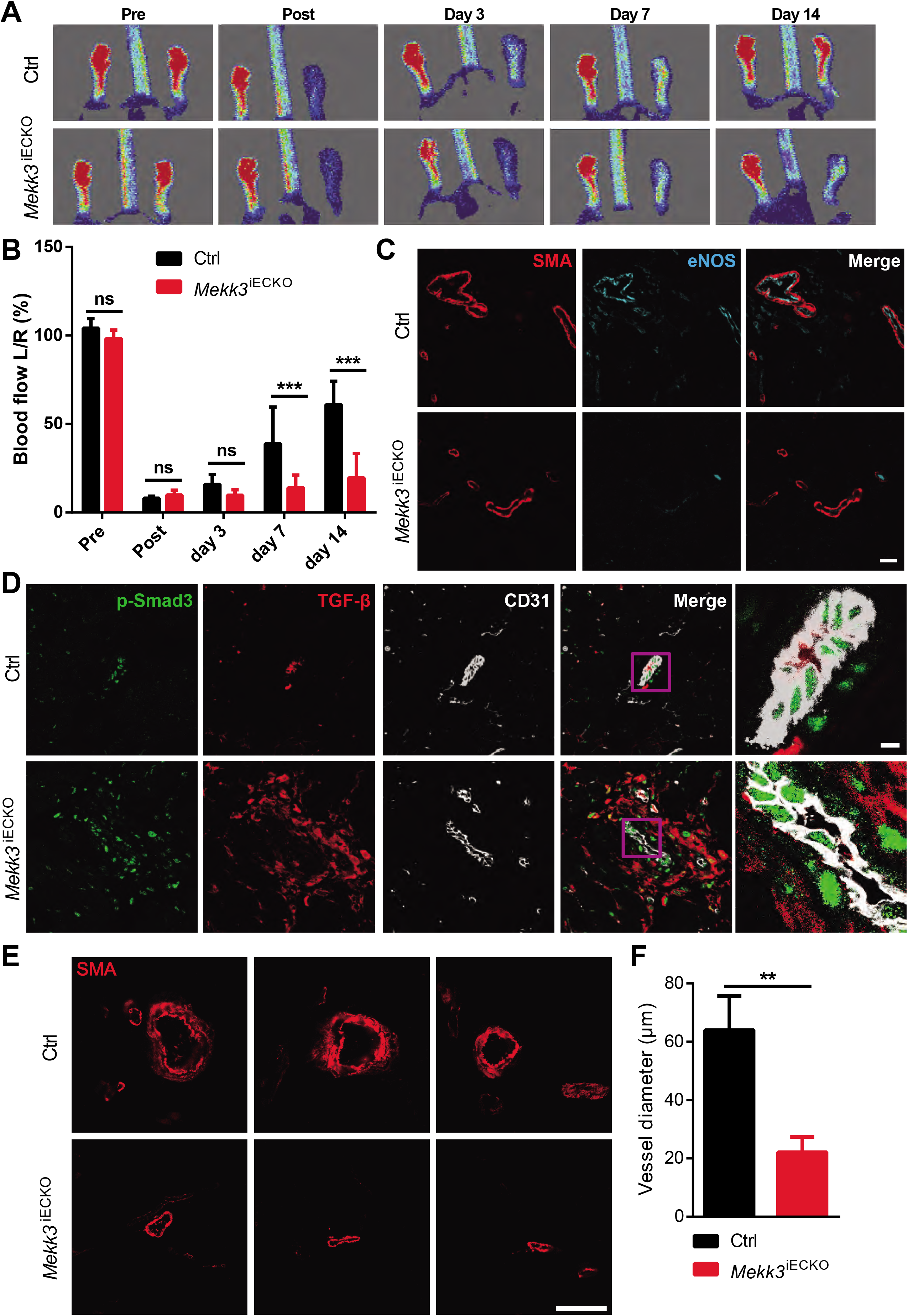
Loss of MEKK3 in ECs impairs blood flow recovery and increases TGFβ signaling in hindlimb ischemia (HLI) model. (**A**) Representative images of blood flow recovery from Ctrl and *Mekk3*^iECKO^ mice at indicated days. (**B**) Quantification of blood flow recovery for Ctrl and *Mekk3*^iECKO^ mice. Data represent mean ± SD, n=6 male mice per group. (**C-D**) Immunostaining of eNOS and SMA (**C**), TGFβ and p-Smad3 (**D**) in gracilis muscle blood vessels from Ctrl and *Mekk3*^iECKO^ mice at day 7. Scale bar: 20μm. (**E-F**) SMA immunostaining and quantification of blood vessels diameter for Ctrl and *Mekk3*^iECKO^ mice. Data represent mean ± SEM, n=6 male mice per group. Scale bar: 50μm. \*\**P* < 0.01, \*\*\**P* < 0.001, ns: not significant, calculated by unpaired *t*-test (F), and two-way ANOVA with Sidak’s multiple comparison tests (B).

To further evaluate a potential contribution of EndMT to observed changes in the vasculature, we carried out immunohistochemical analysis of lung sections from control and *Mekk3*^iECKO^ mice and observed a significant increase in TGFβR1, TGFβ and phospho-Smad3 and p-Smad2 expression (Fig. 5 A-D). To demonstrate the presence of EndMT and assess its contribution to vascular remodeling, we crossed *Mekk3*^iECKO^ mice onto the mTmG background, generating a *Cdh5CreER^*T2*^;Mekk3^flfl^;mTmG* strain. *Mekk3* deletion was induced at 6 weeks of age. Four weeks later mice were sacrificed, and their vasculature examined. A number of pulmonary vessels demonstrated the presence of SMA+ endothelial-derived cells in the expanding neointima and capillaries undergoing arterialization, confirming the presence of EndMT (Fig. 5E and Fig. S7A). Similarly, examination of peripheral tissue demonstrated the presence of endothelial-derived smooth muscle cells in the subintimal of vessels in the kidney, heart and liver (Fig. 5 F-H and Fig. S7 B and C).

**Figure 5.**
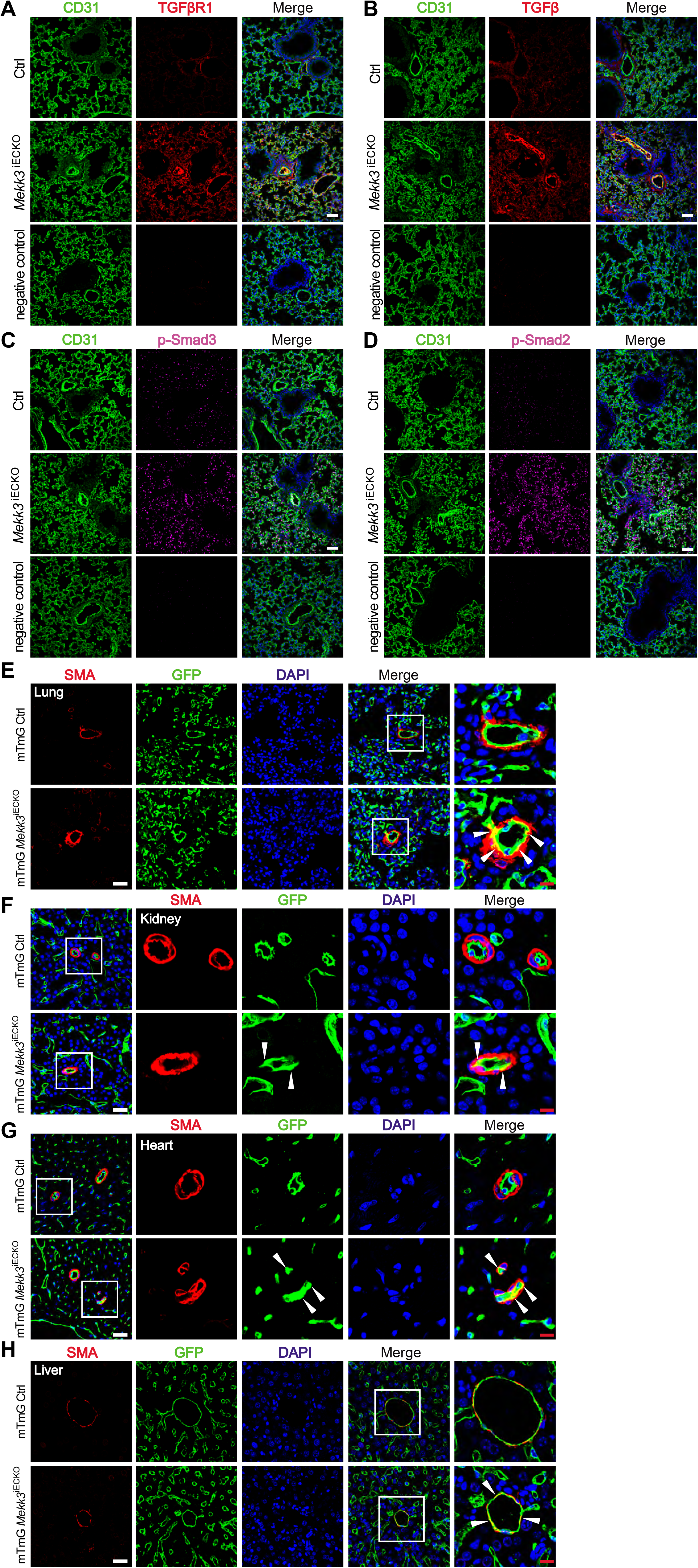
Loss of MEKK3 in the endothelium induces EndMT. (**A-D**) Representative immunostaining of TGFβR1 (**A**), TGFβ (**B**), p-Smad3 Ser423/425 (**C**) and p-Smad2 Ser465/467 (**D**) in lung sections from Ctrl and *Mekk3*^iECKO^ mice at 4 weeks after tamoxifen injection. Scale bar: 50μm. (**E-G**) Representative GFP, SMA and DAPI staining in lung (**E**), kidney (**F**), heart (**G**) and liver (**H**) from mTmG Ctrl and mTmG *Mekk3*^iECKO^ mice at 4 weeks after tamoxifen injection. Scale bar: 25μm (white), 8μm (red). Arrowheads point to endothelial cells expressing SMA.

Prior studies linked EndMT and decreased eNOS to growth of atherosclerotic plaques (24–27). To examine the effect of endothelial MEKK3 deletion on atherosclerosis, *Mekk3*^iECKO^ mice were crossed onto the *Apoe*^−/−^ background, after induction of the *Mekk3* gene deletion at 6 weeks of age, placed on high cholesterol/high fat diet (HCHFD) (Fig. 6A). When sacrificed 8 weeks later, *Apoe*^−/−^;*Mekk3*^iECKO^ mice exhibited much larger atherosclerotic plaques with larger necrotic core, a key characteristic of unstable and vulnerable plaques, both in the aortic root and brachycephalic arteries compared to *Apoe*^−/−^ controls (Fig. 6 B-E), and extensive macrophage invasion of atherosclerotic plaques (Fig. S8).

**Figure 6.**
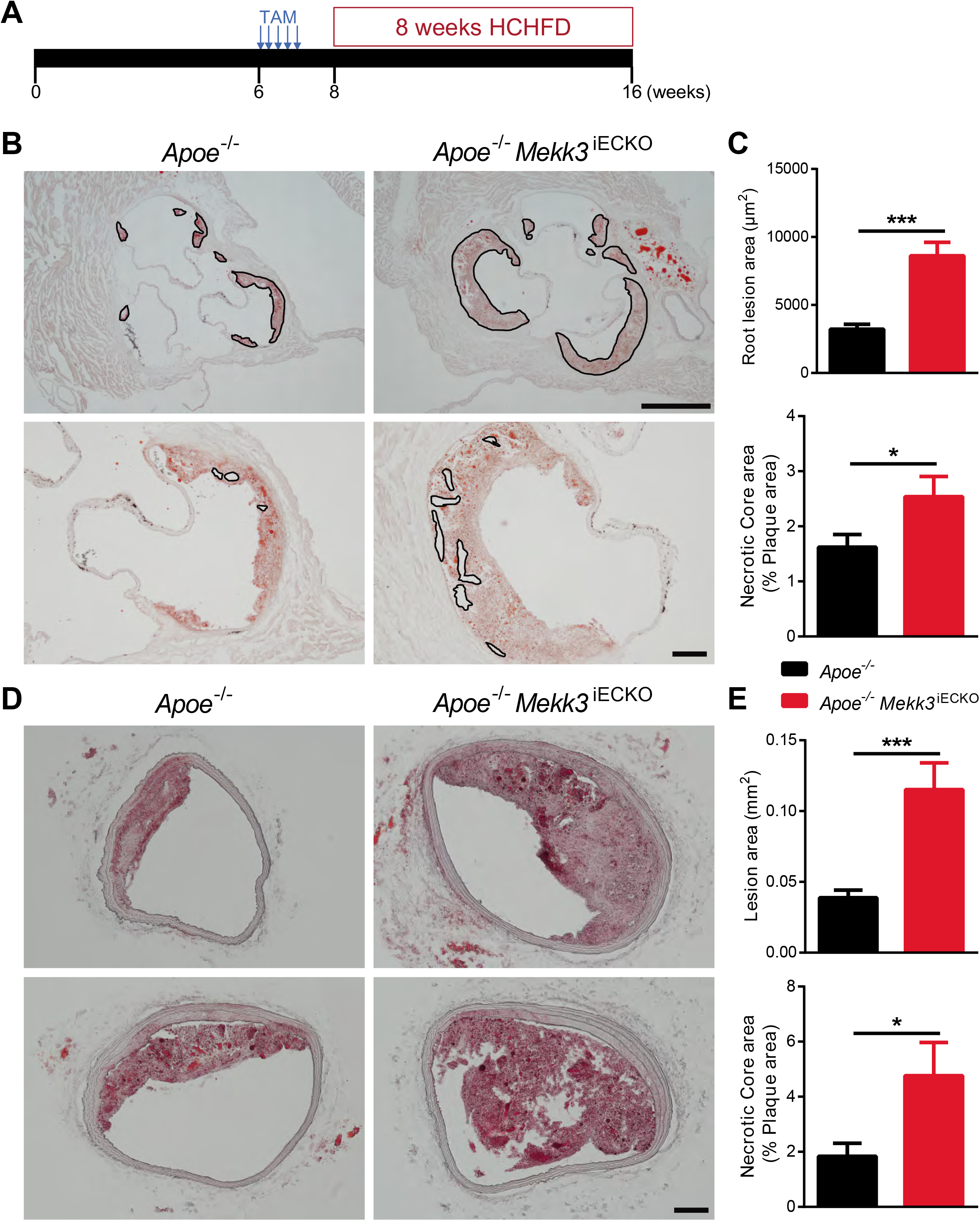
Endothelial MEKK3 knockout increases atherosclerotic plaque growth. (**A**) Experiment timeline: at 6 weeks, *Apoe*^−/−^ mice and *Apoe*^−/−^ *Mekk3*^iECKO^ mice were intraperitoneally injected tamoxifen for 5 consecutive days. From 8 weeks, mice were fed HCHFD for 8 weeks, then were sacrificed for atherosclerosis analysis. (**B**) Representative Oil-Red-O (ORO) staining of aortic root. Scale bar: 100μm. (**C**) Quantification of aortic root lesion area and necrotic core area for *Apoe*^−/−^ mice (n=10 male mice) and *Apoe*^−/−^ *Mekk3*^iECKO^ mice (n=7 male mice). (**D**) Representative ORO staining of brachiocephalic artery. Scale bar: 100μm. (**E**) Quantification of brachiocephalic artery lesion area and necrotic core area for *Apoe*^−/−^ mice (n=10 male mice) and *Apoe*^−/−^ *Mekk3*^iECKO^ mice (n=7 male mice). Data represent mean ± SEM. \**P* < 0.05, \*\*\**P* < 0.001, calculated by unpaired *t*-test.

### Genetic deletion of endothelial TGFβR1 attenuated MEKK3 deficiency-induced hypertension

To test if the observed induction of inward remodeling and EndMT after endothelial MEKK3 knockout are indeed due to activation of TGFβ signaling, HUVECs, following knockdown of MEKK3 expression, were treated either with a TGFβR inhibitor SB431542 or a siRNA-mediated knockdown of TGFβR1 and R2. Both treatments resulted in significant reduction of expression of EndMT markers SM22a, Cdh2 (N-cadherin) and fibronectin (Fig. S9 A-C). To verify these observations in vivo, we combined endothelial MEKK3 knockout with endothelial-specific TGFβR1 knockout by crossing *Cdh5-CreER^*T2*^ Mekk3*^fl/fl^ and *Tgfbr1*^fl/fl^ mice. The generated mouse line, *Cdh5CreER^*T2*^*; *Mekk3*^fl/fl^;*Tgfbr1*^fl/fl^, was subjected to tamoxifen induction at 6 weeks of age (*Mekk3;Tgfbr1* iEC^−/−^, hereafter referred to as DKO). Compared to *Mekk3*^iECKO^ mice, DKO mice showed significantly lower increase in RV systolic pressure and, correspondingly, a reduced increase in RV mass size (Fig. 7 A and B). Whole lung sections showed a decrease in the number of SMA-positive arterial blood vessels indicating reduced EndMT (Fig. 7C). This was further confirmed by p-Smad3 immunocytochemistry (Fig. 7D). We also checked systemic blood pressure at 8 weeks after tamoxifen induction, and found that DKO mice showed significantly lower increase in LV systolic pressure and LV mass (Fig. 7 E and F), as well as systolic and diastolic blood pressure (Fig. 7 G and H) compared to *Mekk3*^iECKO^ mice. Finally, in agreement with the data, the diameter of femoral artery almost returned to normal (Fig. 7 I and G).

**Figure 7.**
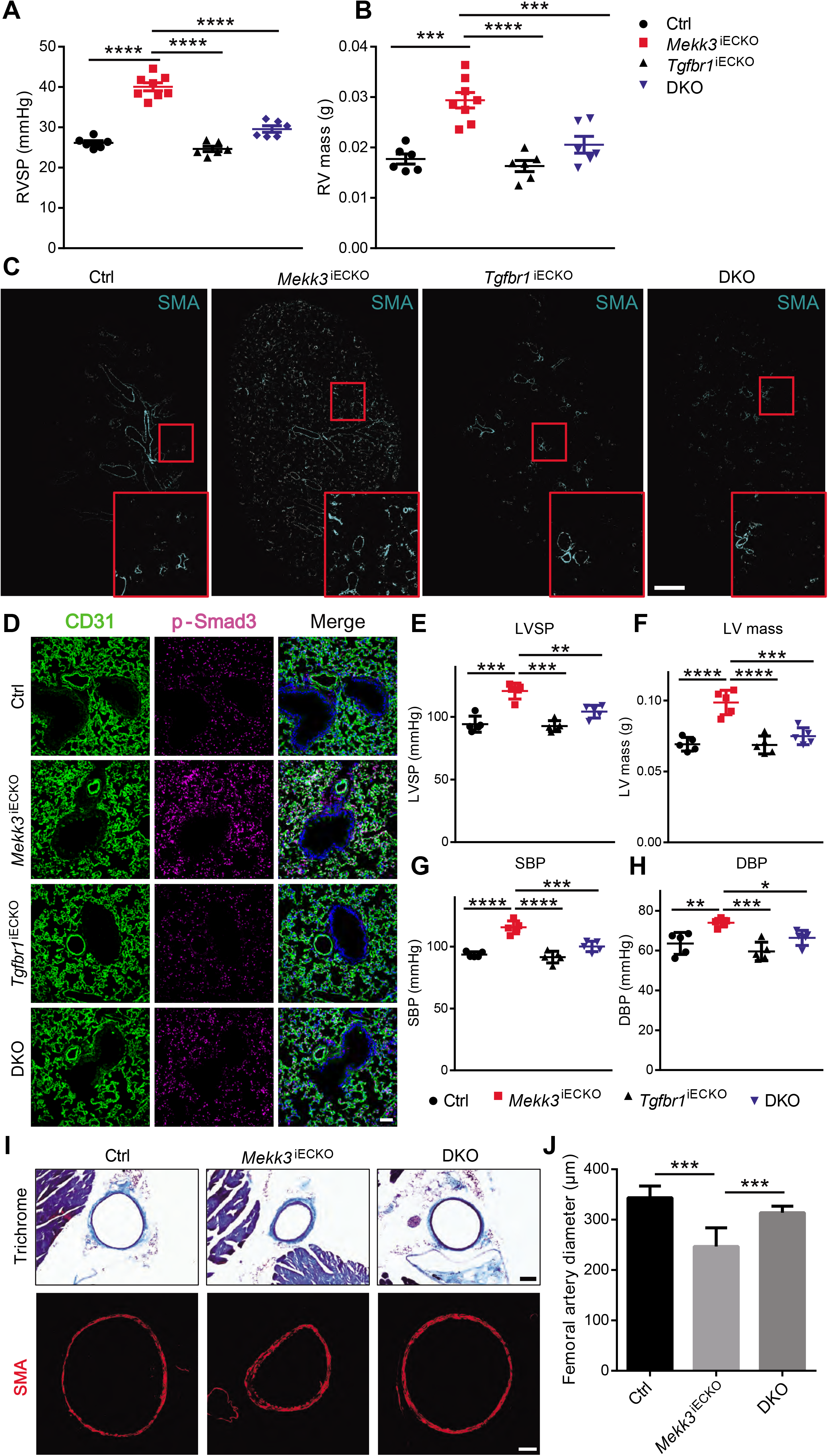
Genetic deletion of endothelial TGFβR1 attenuates MEKK3 deficiency-induced hypertension. (**A-B**) Right ventricle systolic pressure (RVSP) and RV mass of Ctrl (n=6, 3 male and 3 female) and *Mekk3*^iECKO^ (n=8, 4 male and 4 female), *Tgfbr1*^iECKO^ (n=6, 3 male and 3 female), and DKO mice (n=6, 3 male and 3 female) at 3 weeks after tamoxifen injection. (**C**) Representative immunostaining of SMA of Ctrl and *Mekk3*^iECKO^, *Tgfbr1*^iECKO^, and DKO entire lungs. Scale bar: 1mm. (**D**) Immunostaining of p-Smad3 in Ctrl and *Mekk3*^iECKO^, *Tgfbr1*^iECKO^, and DKO lungs. Scale bar: 50μm. (**E-H**) Left ventricle systolic pressure (LVSP), LV mass, Systolic blood pressure (SBP) and Diastolic blood pressure (DBP) of Ctrl and *Mekk3*^iECKO^, *Tgfbr1*^iECKO^, and DKO mice at 8 weeks after tamoxifen injection. (**I**) Representative Masson Trichrome staining (up) and SMA staining (down) of Ctrl, *Mekk3*^iECKO^ and DKO mice femoral artery sections. Scale bar: 100μm (up), 50μm (down). (**J**) Quantification of femoral artery diameter from Trichrome-stained sections (n=30 sections from 3 pairs of mice per group). (**E-H**) n=5 mice (3 male and 2 female) for each group. Data represent mean ± SEM. \**P* < 0.05, \*\**P* < 0.01, \*\*\**P* < 0.001, \*\*\*\**P* < 0.0001, calculated by one-way ANOVA with Tukey’s multiple comparison tests.

## Discussion

The results of this study demonstrate that EC-specific knockout of *Mekk3* in adult mice leads to extensive inward remodeling that affects both systemic and pulmonary vasculatures. At the molecular level, this effect was traced to activation of TGFβ signaling and development of EndMT. These findings, therefore, link MEKK3 to TGFβ signaling and to regulation of inward remodeling.

Vascular remodeling is an important mechanism that allows blood vessels to adapt to blood flow demands. Remodeling is critical to both vascular development and adaptation to disease processes such hypertension or atherosclerosis (1). Sustained changes in blood flow are thought to be critical to both outward and inward remodeling, with the former linked to increased flow and the latter decreased flow (1, 4). Outward remodeling is generally attributed to increased flow and is reasonably well understood. Inward remodeling, a process that leads to first functional and then permanent reduction in arterial diameter, is a particularly vexing problem as it plays a critical role in disease progression in such common illnesses as essential and pulmonary hypertensions, as well as in disease states associated with vascular wall inflammation and reduced flow states. Despite its functional and pathogenic importance, molecular mechanisms controlling inward remodeling are poorly understood.

In this study, we have demonstrated for the first time that deletion of MEKK3 in ECs in adult mice induces inward remodeling in both systemic and pulmonary vessels that leads, over time, to the development of hypertension in both vascular beds. In hyperlipidemic mice endothelial MEKK3 deletion was associated with increased inward remodeling and accelerated atherosclerosis. The key driver in all of these cases was induction of endothelial TGFβ signaling due to an increase in endothelial expression of TGFβ receptors and ligands. Similar to other causes of sustained TGFβ signaling, this led to the appearance of EndMT, both in vitro and in vivo.

In ECs, MEKK3 phosphorylates and activates MEK5, which phosphorylates and activates ERK5, which then mediates induction of the transcription factors KLF2 and 4 (28, 29). This pathway is strongly activated by FSS and mediates induction of many genes responsible for the anti-inflammatory, stabilizing effects of physiological FSS. In CCMs, MEKK3 is also activated and KLF2/4 induced, but in this case by a combination of CCM2 deletion and inflammatory factors, which, combined, drives the disease process (8, 30). In the current study, MEKK3 deletion induced arterial inward remodeling. We also reported FSS activates Smad2/3 signaling specifically at low shear levels with a return to baseline at physiological shear, which drives artery inward remodeling (31). That study also found that MEKK3/KLF2 was responsible for inhibiting Smad2/3 at physiological shear stress, thus shaping the dose-response curve so that Smad2/3 was maximal at low shear. The current findings elaborate a fundamental mechanism in fluid shear stress homeostasis (32). Shear stress at physiological levels activates the MEKK3-KLF2/4 pathway to promote vascular stabilization. When flow decreases, MEKK3 activity and KLF2/4 expression decrease, Smad2/3 activity increases, and vessels undergo inward remodeling. The resultant decrease in lumen diameter restores shear stress to its physiological level, MEKK3 and Smad2/3 return to their previous levels and vessel stability is restored.

In contrast to a transient reduction in FSS, endothelial MEKK3 knockout leads to strong and sustained loss of MEKK3/KLF2 and activation of Smad2/3. Fate-mapping studies demonstrated the appearance of endothelial-derived smooth muscle cells in distal pulmonary arteries, a process associated with distal arterial bed muscularization and the development of pulmonary arterial hypertension (33, 34). In this regard, it is interesting to note a report of an association between a germline heterozygous missense mutation in KLF2 exon 2, c.862C>T p.H288Y and familial pulmonary arterial hypertension (35). Furthermore, deletion of endothelial TGFβR1 abolished both the development of EndMT and pulmonary hypertension. These results strongly implicate the MEKK3/KLF2 axis in pathogenesis of pulmonary arterial hypertension.

Atherosclerosis is another disease process where EndMT has been identified as an important driver of pathogenesis (24, 25, 36, 37). Endothelial MEKK3 deletion in *Apoe*^−/−^ mice showed a significant increase in atherosclerotic plaque formation and necrotic core size. Taken together with pulmonary hypertension data, these results provide a clear functional link between MEKK3 deletion, induction of TGFβ signaling, the onset of EndMT, and the development of EndMT-induced pathogenic events.

While there is a clear connection between EndMT and pulmonary hypertension and there are reports of its association with portal hypertension (38, 39), a relationship between EndMT and systemic hypertension has never being reported. The increase in systemic blood pressure in *Mekk3*^iECKO^ mice paralleled that in pulmonary arterial pressure and both sets of vessels showed a similar extent of inward remodeling. As with the pulmonary vasculature, there was a clear evidence of inward remodeling in systemic arteries that linked contributed to hypertension. Thus, EndMT-driven inward remodeling of peripheral arteries may have well contributed to the development of systemic hypertension.

The development of EndMT may not be the only factor responsible for the appearance of systemic hypertension. We also observed a decrease in NOS3 (eNOS) expression that almost certainly led to a reduction in NO production. There also was an increase in plasma levels of a vasoconstricting hormone endothelin-1. Both of these findings are consistent with a known ERK5-KLF2 and ERK1/2-dependence of eNOS expression (40–43) and endothelin-1 production (23). In this regard it is interesting to speculate that chronic vascular inflammation described in association with systemic hypertension (44, 45), may play a role in the development of EndMT as we have recently proposed (46).

The increase in TGFβ signaling induced by MEKK3 knockout is likely driven by a decrease in expression of let-7 miRNAs family members. ECs normally have high level of let-7 isoforms expression that is maintained by continued FGF signaling input (21). A reduction in FGF signaling, for example deletion of FGFR1, results in a dramatic fall in let-7 expression and increased expression of TGFβ receptor and ligands. This is also the reason for the onset of EndMT under atherosclerotic conditions as chronic vascular inflammation leads to the loss of FGFR1 expression (24, 46). In agreement with these results, endothelial MEKK3 knockout resulted in a significant decrease in let-7 miRNAs levels, likely due to the observed reduction in ERK1/2 signaling. While MEKK3 regulates both ERK1/2 and ERK5 signaling, the observed effect on let-7 I is likely mediated by ERK1/2 given a prior study that demonstrated a clear link between endothelial ERK1/2 deletion, let-7 levels and activation of TGFβ signaling (23).

Given the importance of MEKK3 in adult ECs uncovered here, it is interesting to speculate how alterations in normal regulation of its expression and activity may lead to the loss of its protective function and initiation of pathogenesis. One potential factor, with already established connection to vascular remodeling, is shear stress. But how shear regulates MEKK3 activity is unknown. Two molecular regulators of MEKK3 have been identified. MEKK3 binds Gab1 in a Shp2-dependent manner, leading to a decrease in its activity (47). MEKK3 also binds to and is inhibited by CCM2 (9, 10). As both cell-cell junctions and Shp2 are implicated in FSS signaling, flow could conceivably activate MEKK3 by reversing one or more of these inhibitory interactions. Further work will be required to test these hypotheses and determine molecular mechanisms.

How vessel pathology disrupts or overcomes these pathways is of critical importance. A likely mechanism involves endothelial inflammatory activation, which occurs in nearly all vessel pathologies. Inflammatory cytokines sensitize ECs to TGFβ, thus activating Smad2/3 signaling independent of flow (48). We also identified activation of Smad2/3 by low FSS as the critical signal for homeostatic inward remodeling (31). Thus, by activating Smad2/3, chronic inflammation may mimic these signals to trigger reduction of vascular lumen size. In non-ECs, MEKK3 plays a critical role in mediating pro-inflammatory signals and Toll-like receptor (TLR) signaling (49, 50). MEKK3 deletion in ECs may also change the course of vascular inflammation in response to the circulating pro-inflammatory factors systemically and possibly during the development of atherosclerosis.

In summary, MEKK3 plays an important role in maintenance of endothelial normalcy in the adult vasculature. Its knockout in adult ECs leads to activation of TGFβ signaling resulting in EndMT. This, in turn, induces inward remodeling eventually resulting in the development of both pulmonary and systemic hypertension.

## Materials and Methods

### Mice and genotyping

*Mekk3^fl^*^/*fl*^ mice, *Cdh5-CreER*^*T2*^ mice and *mTmG* reporter mice have been previously described (10). *Mekk2*^−/−^ mice have been reported before (20). *Apoe*^−/−^ mice were purchased from Jackson laboratory (Stock No. 002052). *Tgfbr1^fl^*^/*fl*^ mice are provided by Dr. Anne Eichmann laboratory (Yale University). All mice are C57BL/6 background, and both male and female mice were used for experiments in this study. To induce gene deletion in adults, mice were intraperitoneally injected with 1.5mg tamoxifen (Sigma) for five consecutive days. All mouse protocols and experimental procedures were approved by Yale University Institutional Animal Care and Use Committee (IACUC).

#### Genotyping was performed using the following PCR primers

*Mekk3^fl^*^/*fl*^ (5’-ATG TGA AGC TTG GGG ATT TTG-3’, and 5’-TGG TTA GAC TCA CTG GTC AGA GAC-3’); *Cdh5-CreER*^*T2*^ (5’-CCA AAA TTT GCC TGC ATT ACC GGT CGA TGC-3’, and 5’-ATC CAG GTT ACG GAT ATA GT-3’); *mTmG* (5’-CTC TGC TGC CTC CTG GCT TCT-3’, 5’-CGA GGC GGA TCA CAA GCA ATA-3’, and 5’-TCA ATG GGC GGG GGT CGT T-3’); *Tgfbr1^fl^*^/*fl*^ (5’-ACC CTC TCA CTC TTC CTG AGT-3’, 5’-ATG AGT TAT TAG AAG TTG TTT-3’, and 5’-GGA ACT GGG AAA GGA GAT AAC-3’).

### RV and LV hemodynamic measurements

RVSP was measured with a 1.4F pressure transducer catheter (Millar Instruments) and LabChart software (ADInstruments). Briefly, mice were anesthetized with 2% isoflurane, then the catheter was inserted through the right jugular vein into the right ventricle. For SBP and DBP, the catheter was inserted into the carotid artery. For LVSP, the catheter was inserted through the carotid artery into the left ventricle. RVSP, LVSP, SBP and DBP were all recorded and analyzed with LabChart software (ADInstruments).

### Micro-CT Angiography

For microcomputed tomography (micro-CT) of the lungs, mice were anesthetized with 2% isoflurane, a cannula was inserted into an exposed trachea and connected to a ventilator (120 strokes per min). Then the thorax was opened and the main pulmonary artery instrumented with a flared PE160 catheter that was secured with a 6-0 silk suture. Normal saline was used to perfuse the lungs for 8-10 min followed by perfusion with 10% formaldehyde for 20min. 10% Bismuth nanoparticles in 5% gelatin (CT contrast agent) were then injected into the lung through the PE160 catheter after flush and fixation. Then the lung was covered by ice water-soaked gauze to solid the contrast agent. The lungs were scanned with a specimen micro-CT imaging scanner (GE healthcare) with a cone beam-filtered back projection algorithm, set to a 0.014mm effective detector pixel size. Micro-CT acquisition parameters were 80-kVp x-ray tube voltage, 80-mA tube current, 3000 millisecond per frame exposure time, 1×1 detector binning model, 360 and 0.4 increments per view. For micro-CT of the aorta and hindlimb vasculature (including femoral and tibia artery), euthanized mice were flushed with saline, perfused with 10% formaldehyde at physiologic pressure, and injected with 0.7ml contrast agent in the descending aorta. Then the mice were immediately chilled in ice, and dissected aorta and hindlimb were fixed in 2% formaldehyde overnight. The images were acquired with a specimen micro-CT imaging scanner (GE healthcare), using a cone beam filtered back projection algorithm, set to an 8-27μm thickness. The mCT data were processed using real time 3D volume rendering software (version 3.1, Vital Images, Inc) and Microview (version 1.15, GE medical system) software to reconstruct three 2D maximum intensity projection images from the raw data. Quantification was performed using a modified Image ProPlus 5.0 algorithm (Media Cybernetics).

### Hind limb ischemia

Briefly, surgical procedures were performed in mice under anesthesia condition. A vertical longitudinal incision was made in the right hind limb (10 mm long). The right common femoral artery and its side branches were dissected and ligated with 6-0 silk sutures spaced 5 mm apart, and the arterial segment between the ligatures was excised. Assessment of tissue perfusion by Laser-Doppler flow-Imaging (LDI) was done by scanning rear paws with the LDI analyzer (Moor Infrared Laser Doppler Imager Instrument, Wilmington, Delaware) before and after the surgical procedure. Low or no perfusion is displayed as dark blue, whereas the highest degree of perfusion is displayed as red. The images were quantitatively converted into histograms with Moor LDI processing software V3.09. Data are reported as the ratio of flow in the right/left (R/L) hind limb and calf regions. Measurement of blood flow was done before and immediately after the surgical procedure and then at days 3, 7, and 14.

### Immunohistochemistry

Samples were fixed in 4% paraformaldehyde (Electron Microscopy Sciences) overnight at 4°C and incubated with 30% sucrose (Sigma) solution in PBS overnight at 4°C. Then samples were embedded in OCT medium (SAKURA) and 8-10μm sections were cut in a cryostat (Leica). For immunohistochemistry, samples were blocked in a blocking buffer (5% donkey serum, 0.2% BSA, 0.3% Triton X-100 in PBS), followed by incubation with primary and secondary antibodies diluted in a blocking buffer. Images were taken using the SP8 confocal microscope (Leica).

### Histological analysis of atherosclerotic lesions

Mice were fed with HCHFD (Research Diets, product no. D12108) for 8 weeks, then were euthanized and perfusion-fixed with PBS and 4% paraformaldehyde via the left ventricle. Mouse hearts and brachiocephalic artery were isolated, fixed in 4% PFA O/N, then dehydrated with 30% sucrose O/N, embedded in OCT, and placed at −80°C. Aortic root and brachiocephalic artery blocks were sectioned using a cryostat. For morphometric analysis, serial sections were cut at 8μm thickness. Every third slide from the serial sections was stained with Oil-Red-O (ORO). Aortic root lesion area was quantified by averaging the lesion areas in 6 sections from the same mouse. Brachiocephalic artery lesion area was quantified by averaging the lesion areas in 9 sections from the same mouse. Lesion areas were quantified with ImageJ (National Institute of Health) software.

### Cell culture and siRNA transfection

HUVECs (Human Umbilical Vein Endothelial Cells) were obtained from Yale Vascular Biology and Therapeutics tissue-culture core laboratory at Passage 1. HPAECs (Human Pulmonary Artery Endothelial Cells) were purchased from Lonza. Both cells were maintained in EMG2 Endothelial Cells Growth Media (Lonza) and used for experiments between P2 and P5. SiRNA transfection was performed with Opti-MEM medium (ThermFisher) and Lipofectamine RNAiMAX (Invitrogen). At day 4 after transfection, cells were harvested for protein and RNA analysis. MEKK3 siRNA were purchased from QIAGEN (Hs_MAP3K3_5, target sequence CAGGAATACTCAGATCGGGAA). TGFβR1 siRNA and TGFβR2 siRNA were from Dharmacon. TGFβR inhibitor SB431542 was purchased from Sigma (S4317).

### RNA isolation and quantitative real-time PCR

RNA was extracted from cells with RNeasy Plus Mini Kit (QIAGEN) according to the manufacturer’s instructions, and the reverse transcription reaction was performed with the iScript Reverse Transcription Supermix for RT-qPCR (BIO-RAD). Then cDNA was amplified by real-time PCR with iQ SYBR Green Supermix (BIO-RAD). The expression of target genes was normalized to expression of housekeeping gene *GAPDH*. Primers for qPCR were listed in Supplemental Table 1.

### RNA-Sequencing Analysis

Total RNA was extracted from HUVECs treated with Ctrl and MEKK3 siRNA (four samples for each group) and quantitated by NanoDrop, and RNA integrity number value was measured with an Agilent Bioanalyzer. Samples were subjected to RNA-sequencing using Illumina NextSeq 500 sequencer (75bp paired end reads). The base calling data from sequencer were transferred into FASTQ files using bcl2fastq2 conversion software (version 2.20, Illumina). The raw reads were aligned to the human reference genome GRCh38 using HISAT2 (version 2.1.0) alignment software. The raw reads were processed using HTSeq (version 0.11.1) to generate read counts for every gene. DESeq2 (version 1.24, using default parameter) was used to pre-process raw data to remove the noise, normalize each sample to correct the batch effect, perform PCA for observing the similarity of replicates, and identify differential expression genes. P values obtained from multiple tests were adjusted using Benjamini-Hochberg correction.

### MicroRNA Real-Time PCR Analysis

Total RNA was extracted from cells with miRNeasy Mini Kit (QIAGEN) according to the manufacturer’s instructions. Reverse transcription reaction was performed with the miScript Reverse Transcription kit (QIAGEN), and cDNA was amplified by real-time PCR with a miScript SYBR Green kit (QIAGEN). Individual miRNA expression was normalized in relation to expression of small nuclear SNORD47. PCR amplification consisted of 10 min of an initial denaturation step at 95°C, followed by 45 cycles of PCR at 95°C for 30s, 60°C for 30s. Let-7 miRNA primers were purchased from QIAGEN.

### Western Blotting

Cells were harvested and lysed for 30 min on ice in RIPA buffer (Roche) containing complete mini protease inhibitors (Roche) and phosphatase inhibitors (Roche). Centrifuge at 14000 rpm for 10 min at 4°C, transfer cell supernatant to new 1.5 ml EP tubes, and boiled with 4X loading buffer (250mM Tris-HCl pH 6.8, 8% SDS, 40% glycerol, 20% β-mercaptoethanol, 0.008% bromophenol blue) at 99°C for 5 min. Cell lysates were resolved by a 4-15% CriterionTM TGXTM Precast Gels (BIO-RAD) SDS–PAGE electrophoresis, and blotted onto a polyvinylidene difluoride (PVDF) membrane (Millipore). The blotted membranes were blocked with 5% non-fat milk and then incubated with specific antibodies diluted in 5% BSA using a standard immunoblotting procedure and detection by ECL (Millipore). ImageJ was used for densitometry quantification of western blot bands.

### Antibodies

We used the following antibodies for immunohistochemistry (IHC) and immunoblotting (IB): Rat anti-Mouse CD31 (BD 550274; IHC 1:200), GAPDH (Cell Signaling 5174S; IB 1:1000), MEKK3 (R&D Systems MAB6095; IB 1:1000), MEKK2 (Abcam ab33918; IB 1:5000), Rat anti-MECA32 (Developmental Studies Hybridoma Bank; IHC 1:15), Fibronectin (BD 610078; IB 1:10000, IHC 1:400), eNOS (BD 610297; IHC 1:400), TGFβR1 (Santa Cruz sc-398; IB 1:1000), TGFβR2 (Santa Cruz sc-400; IB 1:1000), TGFβ (R&D Systems MAB1835; IHC 1:400), p-Smad2 Ser465/467 (Millipore AB3849-I; IHC 1:200), p-Smad3 Ser423/425 (Abcam ab52903; IHC 1:200), SM22 alpha (Abcam ab14106; IB 1:5000, IHC 1:400), α-Smooth Muscle Actin (Sigma A2547; IB 1:1000, IHC 1:200), FGFR1 (Cell Signaling 9740; IB 1:1000), GFP (aves GFP-1020; IHC 1:400), p-ERK1/2 (Cell Signaling 4370; IB 1:1000).

### Mouse lung ECs, aortic smooth muscle cells, cardiomyocytes isolation

Mouse lungs were collected and digested in a solution of 2mg/ml collagenase (Sigma). Cell suspension were then filtered through 70μm sterile cell strainer (Falcon). ECs were isolated using magnetic beads anti-rat IgG (Invitrogen) coated with rat anti-mouse CD31 antibody (BD Biosciences). Aortas were collected and digested in 175U/ml collagenase and 1.25U/ml elastase in HBSS for 15min at RT. Adventitia layer was stripped off. Media layer was minced and further digested with 400U/ml collagenase and 2.5U/ml elastase in HBSS at 37°C for 1h. ECs were selected by anti-mouse CD31 antibody and discarded, and the remaining SMCs were lysed. Hearts were excised and perfused at 37°C with a Langendorff apparatus (ADInstruments) with a Ca^2+^-free Krebs-Henseleit buffer (0.6mM KH_2_PO_4_, 0.6mM Na_2_HPO_4_, 1.7mM MgSO_4_, 4.6mM NaHCO_3_, 10mM HEPES, 14.7mM KCl, 120.3mM NaCl, 30mM taurine, 10mM glucose, 15mM 2,3-butanedione monoxime, pH7.3). Then hearts were perfused and digested with perfusion buffer plus 0.067 mg/ml Liberase Blendzyme 4 (Roche) for 30min. After complete digestion, hearts were minced and filtered. Cells were lysed for RNA extraction with PicoPure RNA isolation kit (Applied Biosystems) according to the manufacturer’s instructions.

### Enzyme-linked immunosorbent assay (ELISA)

Endothelin-1 ELISA was performed on plasma using Endothelin-1 Immunoassay kit (R&D Systems) according to manufacturer’s instructions. Optical density was determined using a microplate reader (BioTek).

### Statistical analysis

Statistical analysis was performed using GraphPad Prism software (GraphPad software Inc.). Specific statistical analysis was clearly stated in each figure legend. A *P* value less than 0.05 was considered significant (\**P* < 0.05, \*\**P* < 0.01, \*\*\**P* < 0.001, \*\*\*\**P* < 0.0001).

## Author contributions

H.D. performed most of the experiments, analyzed data, and prepared figures and manuscript. X.H. measured pressure. Z.W.Z. performed micro-CT. Y.X. and A.N. helped with tissue staining. Y.C. and Y.W. helped with RNA-seq analysis. M.A.S discussed data and wrote manuscript. B.S. provided the *Mekk3* and *Mekk2* mice strains, analyzed data, discussed results, and edited manuscript. M.S. supervised and supported the project, analyzed data, and wrote the manuscript.

## Acknowledgments

We are grateful to Anne Eichmann for TGFβR1floxed mice, Hyung Chun and Brian G. Coon for helpful discussion, Rita Webber and Laran Coon for maintaining mice colonies used in this study. We also thank Nicolas Ricard, Georgia Zarkada, Dou Liu, Yi Xie and Xinbo Zhang for technical help. This work was supported by NIH Grant R01 HL053793 (MS), R01 HL062289 (MS), and NSFC grants 31930035, 91942311 (BS).

## References

1. Baeyens N, Bandyopadhyay C, Coon BG, Yun S, & Schwartz MA (2016) Endothelial fluid shear stress sensing in vascular health and disease. The Journal of clinical investigation 126(3):821–828.

2. Renna NF, de Las Heras N, & Miatello RM (2013) Pathophysiology of vascular remodeling in hypertension. International journal of hypertension 2013:808353.

3. Castorena-Gonzalez JA, Staiculescu MC, Foote C, & Martinez-Lemus LA (2014) Mechanisms of the inward remodeling process in resistance vessels: is the actin cytoskeleton involved? Microcirculation 21(3):219–229.

4. Ward MR, et al. (2001) Low blood flow after angioplasty augments mechanisms of restenosis: inward vessel remodeling, cell migration, and activity of genes regulating migration. Arteriosclerosis, thrombosis, and vascular biology 21(2):208–213.

5. Peti W & Page R (2013) Molecular basis of MAP kinase regulation. Protein science: a publication of the Protein Society 22(12):1698–1710.

6. Yang J, et al. (2000) Mekk3 is essential for early embryonic cardiovascular development. Nat Genet 24(3):309–313.

7. Cullere X, Plovie E, Bennett PM, MacRae CA, & Mayadas TN (2015) The cerebral cavernous malformation proteins CCM2L and CCM2 prevent the activation of the MAP kinase MEKK3. Proceedings of the National Academy of Sciences of the United States of America 112(46):14284–14289.

8. Zhou Z, et al. (2016) Cerebral cavernous malformations arise from endothelial gain of MEKK3-KLF2/4 signalling. Nature 532(7597):122–126.

9. Zhou Z, et al. (2015) The cerebral cavernous malformation pathway controls cardiac development via regulation of endocardial MEKK3 signaling and KLF expression. Developmental cell 32(2):168–180.

10. Fisher OS, et al. (2015) Structure and vascular function of MEKK3-cerebral cavernous malformations 2 complex. Nature communications 6:7937.

11. Hayashi M, et al. (2004) Targeted deletion of BMK1/ERK5 in adult mice perturbs vascular integrity and leads to endothelial failure. The Journal of clinical investigation 113(8):1138–1148.

12. Srinivasan R, et al. (2009) Erk1 and Erk2 regulate endothelial cell proliferation and migration during mouse embryonic angiogenesis. PloS one 4(12):e8283.

13. Yan C, Takahashi M, Okuda M, Lee JD, & Berk BC (1999) Fluid shear stress stimulates big mitogen-activated protein kinase 1 (BMK1) activity in endothelial cells. Dependence on tyrosine kinases and intracellular calcium. The Journal of biological chemistry 274(1):143–150.

14. Li L, et al. (2008) Fluid shear stress inhibits TNF-mediated JNK activation via MEK5-BMK1 in endothelial cells. Biochemical and biophysical research communications 370(1):159–163.

15. Wu K, Tian S, Zhou H, & Wu Y (2013) Statins protect human endothelial cells from TNF-induced inflammation via ERK5 activation. Biochemical pharmacology 85(12):1753–1760.

16. Komaravolu RK, et al. (2015) Erk5 inhibits endothelial migration via KLF2-dependent down-regulation of PAK1. Cardiovascular research 105(1):86–95.

17. Pi X, Yan C, & Berk BC (2004) Big mitogen-activated protein kinase (BMK1)/ERK5 protects endothelial cells from apoptosis. Circulation research 94(3):362–369.

18. Hayashi M & Lee JD (2004) Role of the BMK1/ERK5 signaling pathway: lessons from knockout mice. Journal of molecular medicine 82(12):800–808.

19. Wang X, Chang X, Facchinetti V, Zhuang Y, & Su B (2009) MEKK3 is essential for lymphopenia-induced T cell proliferation and survival. Journal of immunology 182(6):3597–3608.

20. Guo Z, et al. (2002) Disruption of Mekk2 in mice reveals an unexpected role for MEKK2 in modulating T-cell receptor signal transduction. Molecular and cellular biology 22(16):5761–5768.

21. Chen PY, et al. (2012) FGF regulates TGF-beta signaling and endothelial-to-mesenchymal transition via control of let-7 miRNA expression. Cell reports 2(6):1684–1696.

22. Chen PY, Qin L, Tellides G, & Simons M (2014) Fibroblast growth factor receptor 1 is a key inhibitor of TGFbeta signaling in the endothelium. Science signaling 7(344):ra90.

23. Ricard N, et al. (2019) Endothelial ERK1/2 signaling maintains integrity of the quiescent endothelium. The Journal of experimental medicine 216(8):1874–1890.

24. Chen PY, et al. (2015) Endothelial-to-mesenchymal transition drives atherosclerosis progression. The Journal of clinical investigation 125(12):4514–4528.

25. Chen PY, et al. (2019) Endothelial TGF-beta signalling drives vascular inflammation and atherosclerosis. Nature metabolism 1(9):912–926.

26. Kuhlencordt PJ, et al. (2001) Accelerated atherosclerosis, aortic aneurysm formation, and ischemic heart disease in apolipoprotein E/endothelial nitric oxide synthase double-knockout mice. Circulation 104(4):448–454.

27. Chen J, Kuhlencordt PJ, Astern J, Gyurko R, & Huang PL (2001) Hypertension does not account for the accelerated atherosclerosis and development of aneurysms in male apolipoprotein e/endothelial nitric oxide synthase double knockout mice. Circulation 104(20):2391–2394.

28. Chao TH, Hayashi M, Tapping RI, Kato Y, & Lee JD (1999) MEKK3 directly regulates MEK5 activity as part of the big mitogen-activated protein kinase 1 (BMK1) signaling pathway. The Journal of biological chemistry 274(51):36035–36038.

29. Nakamura K & Johnson GL (2003) PB1 domains of MEKK2 and MEKK3 interact with the MEK5 PB1 domain for activation of the ERK5 pathway. The Journal of biological chemistry 278(39):36989–36992.

30. Cuttano R, et al. (2016) KLF4 is a key determinant in the development and progression of cerebral cavernous malformations. EMBO molecular medicine 8(1):6–24.

31. Deng H, et al. (2021) Activation of Smad2/3 signaling by low fluid shear stress mediates artery inward remodeling. Proceedings of the National Academy of Sciences of the United States of America (in press).

32. Baeyens N, et al. (2015) Vascular remodeling is governed by a VEGFR3-dependent fluid shear stress set point. eLife 4.

33. Kovacic JC, et al. (2019) Endothelial to Mesenchymal Transition in Cardiovascular Disease: JACC State-of-the-Art Review. Journal of the American College of Cardiology 73(2):190–209.

34. Dejana E, Hirschi KK, & Simons M (2017) The molecular basis of endothelial cell plasticity. Nature communications 8:14361.

35. Eichstaedt CA, et al. (2017) First identification of Kruppel-like factor 2 mutation in heritable pulmonary arterial hypertension. Clinical science 131(8):689–698.

36. Evrard SM, et al. (2016) Endothelial to mesenchymal transition is common in atherosclerotic lesions and is associated with plaque instability. Nature communications 7:11853.

37. Moonen JR, et al. (2015) Endothelial-to-mesenchymal transition contributes to fibro-proliferative vascular disease and is modulated by fluid shear stress. Cardiovascular research 108(3):377–386.

38. Sato Y & Nakanuma Y (2013) Role of endothelial-mesenchymal transition in idiopathic portal hypertension. Histology and histopathology 28(2):145–154.

39. Nakanuma Y, Sato Y, & Kiktao A (2009) Pathology and pathogenesis of portal venopathy in idiopathic portal hypertension: Hints from systemic sclerosis. Hepatology research: the official journal of the Japan Society of Hepatology 39(10):1023–1031.

40. Chamorro-Jorganes A, et al. (2016) VEGF-Induced Expression of miR-17-92 Cluster in Endothelial Cells Is Mediated by ERK/ELK1 Activation and Regulates Angiogenesis. Circulation research 118(1):38–47.

41. Chen J, et al. (2005) Akt1 regulates pathological angiogenesis, vascular maturation and permeability in vivo. Nature medicine 11(11):1188–1196.

42. Ren B, et al. (2010) ERK1/2-Akt1 crosstalk regulates arteriogenesis in mice and zebrafish. The Journal of clinical investigation 120(4):1217–1228.

43. SenBanerjee S, et al. (2004) KLF2 Is a novel transcriptional regulator of endothelial proinflammatory activation. The Journal of experimental medicine 199(10):1305–1315.

44. Harrison DG, Marvar PJ, & Titze JM (2012) Vascular inflammatory cells in hypertension. Frontiers in physiology 3:128.

45. McMaster WG, Kirabo A, Madhur MS, & Harrison DG (2015) Inflammation, immunity, and hypertensive end-organ damage. Circulation research 116(6):1022–1033.

46. Schwartz MA, Vestweber D, & Simons M (2018) A unifying concept in vascular health and disease. Science 360(6386):270–271.

47. Lerner-Marmarosh N, et al. (2003) Inhibition of tumor necrosis factor-[alpha]-induced SHP-2 phosphatase activity by shear stress: a mechanism to reduce endothelial inflammation. Arteriosclerosis, thrombosis, and vascular biology 23(10):1775–1781.

48. Min E & Schwartz MA (2019) Translocating transcription factors in fluid shear stress-mediated vascular remodeling and disease. Experimental cell research 376(1):92–97.

49. Yang J, et al. (2001) The essential role of MEKK3 in TNF-induced NF-kappaB activation. Nature immunology 2(7):620–624.

50. Huang Q, et al. (2004) Differential regulation of interleukin 1 receptor and Toll-like receptor signaling by MEKK3. Nature immunology 5(1):98–103.

