## Supplemental figures and table for "MEKK3-TGFβ crosstalk regulates inward arterial remodeling"

[6]Co-first authors, [7]Co-senior authors, [8]Lead contact

Address correspondence to:

Prof. Michael Simons

Yale University School of Medicine

300 George St, Suite 773

New Haven, CT 06511

**Supplemental Figure Legends**

**Figure S1. Loss of MEKK3 in ECs induces cardiac hypertrophy.** (**A**) Q-PCR analysis of *Mekk3* expression in isolated lung ECs, aortic smooth muscle cells and cardiomyocytes from Ctrl and *Mekk3*$^{iECKO}$ mice (n=3). Data represent mean ± SD. ****$P$ < 0.0001, ns: not significant, calculated by unpaired *t*-test. (**B**) Representative higher magnification H&E images of RV and LV for Ctrl and *Mekk3*$^{iECKO}$ mice at 4 weeks after tamoxifen injection. Scale bar: 500µm. (**C**) Representative right ventricle systolic pressure (RVSP) tracing for Ctrl and *Mekk3*$^{iECKO}$ mice at 4 weeks after tamoxifen injection. (**D**) Representative left ventricle systolic pressure (LVSP) tracing for Ctrl and *Mekk3*$^{iECKO}$ mice at 8 weeks after tamoxifen injection. (**E**) Representative systolic blood pressure (SBP) and diastolic blood pressure (DBP) tracing for Ctrl and *Mekk3*$^{iECKO}$ mice at 8 weeks after tamoxifen injection.

**Figure S2. Heart rate and ECG for Ctrl and *Mekk3*$^{iECKO}$ mice.** (**A**) Heart rate (BPM) for Ctrl and *Mekk3*$^{iECKO}$ mice at 2, 4 and 8 weeks after tamoxifen injection. n=5 mice for each group. Data represent mean ± SEM. ns: not significant, calculated by two-way ANOVA with Tukey's multiple comparison tests. (**B**) Representative electrocardiogram (ECG) tracing for Ctrl and *Mekk3*$^{iECKO}$ mice at 4 and 8 weeks after tamoxifen injection.

**Figure S3. *Mekk2* knockout mice don't develop hypertension.** (**A**) Western blot analysis of MEKK2 expression in Ctrl and *Mekk2*$^{-/-}$ lungs. (**B**) RVSP of Ctrl and *Mekk2*$^{-/-}$ mice at 6 months old. (**C**) LVSP of Ctrl and *Mekk2*$^{-/-}$ mice at 6 months old. (**B-C**) n=4 male mice for each group. Data represent mean ± SEM. ns: not significant, calculated by unpaired *t*-test.

**Figure S4. TGFβR2 staining in lungs of *Mekk3*$^{iECKO}$ mice.** (**A**) Representative TGFβR2 staining of Ctrl and *Mekk3*$^{iECKO}$ entire lungs at 4 weeks after tamoxifen injection. Scale bar: 1mm.

**Figure S5. Knockdown of MEKK3 in HPAECs induces EndMT.** (**A**) Q-PCR analysis of EndMT markers expression in human pulmonary artery endothelial cells (HPAECs) treated with Ctrl or MEKK3 siRNA. Data represent mean ± SD. $^{**}P < 0.01$, $^{***}P < 0.001$, calculated by unpaired *t*-test. (**B**) Western blot analysis of EndMT markers expression in HPAECs treated with Ctrl or MEKK3 siRNA. (**C**) Immunostaining of LIN28 in Ctrl and $Mekk3^{iECKO}$ mice lungs. Scale bar: 50μm. Arrowheads point to endothelial cells expressing LIN28.

**Figure S6. Loss of MEKK3 in ECs impairs FGF2-ERK1/2-Let7 signaling pathway.** (**A-B**) ERK1/2 activation upon (**A**) FGF2 (100ng/ml) and (**B**) VEGF165 (50ng/ml) treatment in HUVECs treated with Ctrl or MEKK3 siRNA. (**C-D**) Smad1/5/9 activation upon (**C**) BMP9 (10ng/ml) and (**D**) BMP6 (50ng/ml) treatment in HUVECs treated with Ctrl or MEKK3 siRNA. (**E**) Q-PCR analysis of FGFR1 (n=6) expression in HUVECs treated with Ctrl or MEKK3 siRNA. Data represent mean ± SEM. (**F**) Western blot analysis of FGFR1 expression in HUVECs treated with Ctrl or MEKK3 siRNA. ns: not significant, calculated by unpaired *t*-test.

**Figure S7. Additional images showing EndMT.** (**A-C**) Representative GFP and SMA staining in lung (**A**), kidney (**B**), and liver (**C**) from mTmG Ctrl and mTmG $Mekk3^{iECKO}$ mice at 4 weeks after tamoxifen injection. Scale bar: 50μm. Arrowheads point to endothelial cells expressing SMA.

**Figure S8. F4/80 staining in atherosclerotic plaque.**

(**A**) Representative F4/80 staining in brachiocephalic artery lesion from $Apoe^{-/-}$ mice and $Apoe^{-/-}$ $Mekk3^{iECKO}$ mice. Scale bar: 100μm.

**Figure S9. Suppression of TGFβR signaling rescues MEKK3-knockout-induced EndMT.** (**A**) Q-PCR analysis of MEKK3, SM22α, fibronectin (FN) and N-Cadherin expression in HUVECs treated with Ctrl or MEKK3 siRNA in addition to TGFβR inhibitor. (**B**) Western Blot and (**C**) Q-PCR analysis of EndMT markers expression in HUVECs treated with Ctrl or MEKK3 siRNA in addition to TGFβR1/R2 siRNA. n=3. Data represent mean ± SD. $^{**}P < 0.01$, $^{***}P < 0.001$, ns: not significant, calculated by one-way ANOVA with Tukey's multiple comparison tests.

#### Figure S1

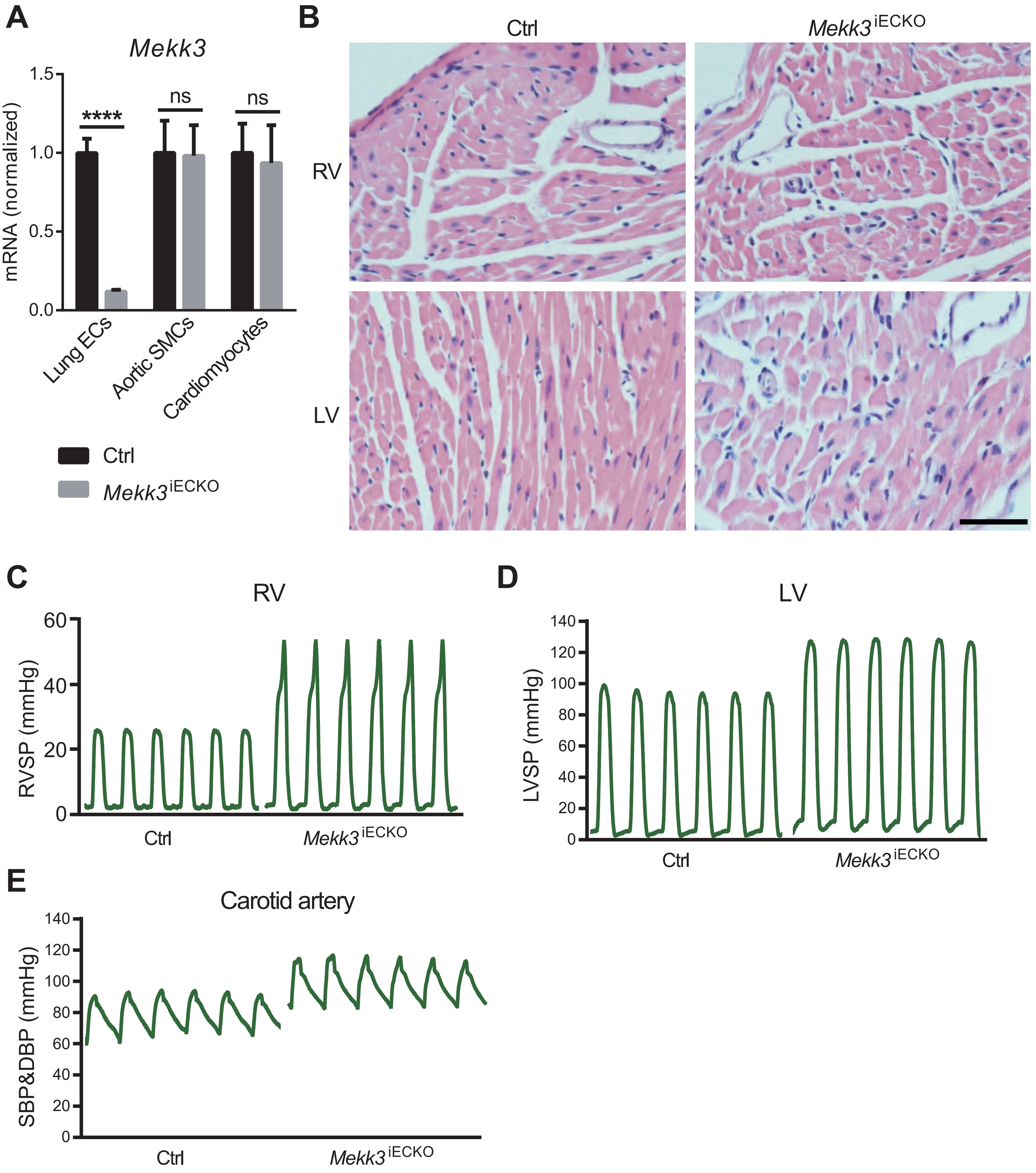

A
Mekk3
mRNA (normalized)
1.5
1.0
0.5
0.0
****
ns
ns
Lung ECs
Aortic SMCs
Cardiomyocytes
Ctrl
Mekk3 iECKO
B
Ctrl
Mekk3 iECKO
RV
LV
C
RV
RVSP (mmHg)
60
40
20
0
Ctrl
Mekk3 iECKO
D
LV
LVSP (mmHg)
140
120
100
80
60
40
20
0
Ctrl
Mekk3 iECKO
E
Carotid artery
SBP&DBP (mmHg)
140
120
100
80
60
40
20
0
Ctrl
Mekk3 iECKO

### Figure S2

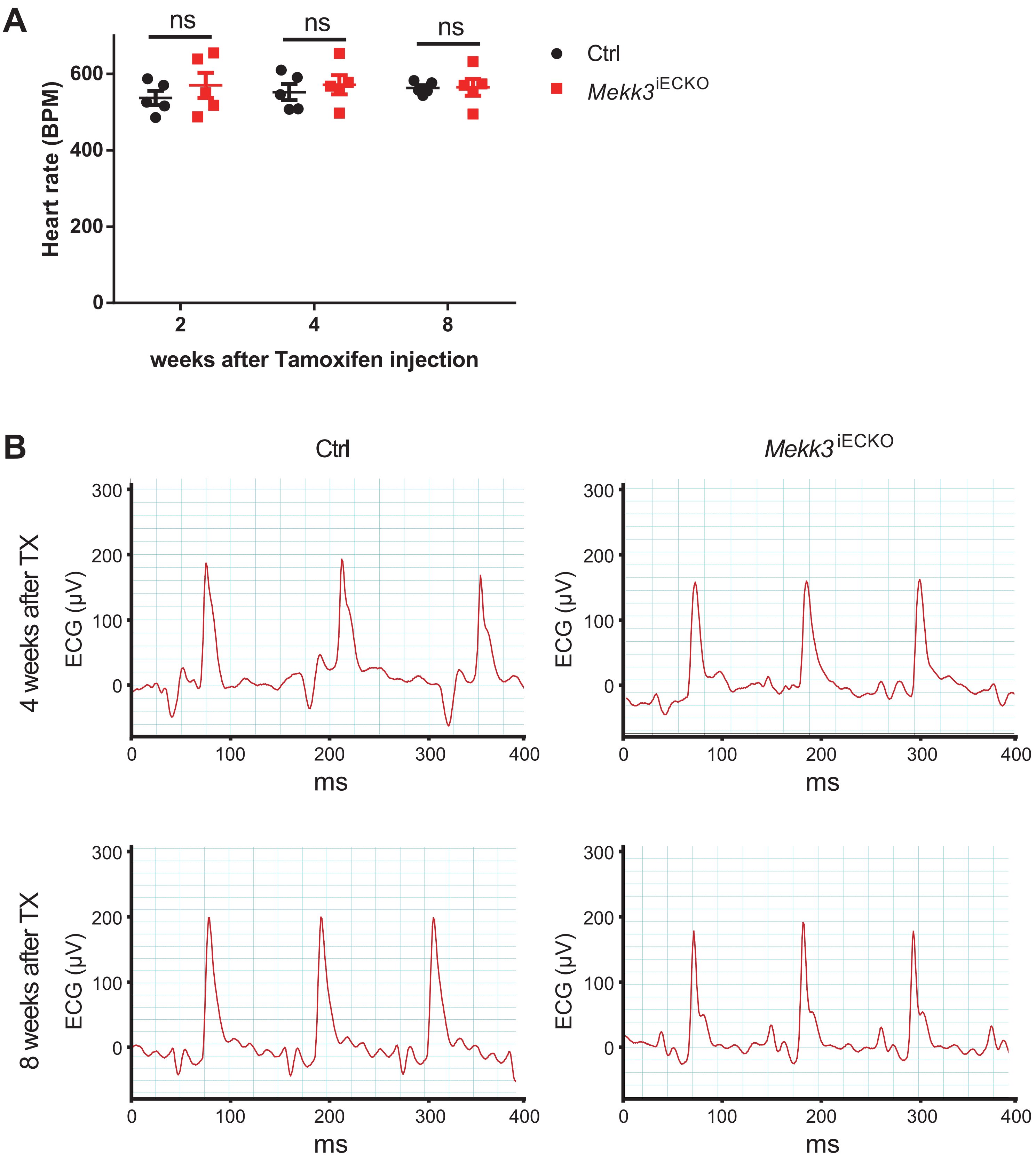
A
ns
ns
ns
Ctrl
Mekk3iECKO
Heart rate (BPM)
600
400
200
0
2
4
8
weeks after Tamoxifen injection
B
Ctrl
Mekk3iECKO
4 weeks after TX
8 weeks after TX
ECG (µV)
300
200
100
0
0
100
200
300
400
ms

### Figure S3

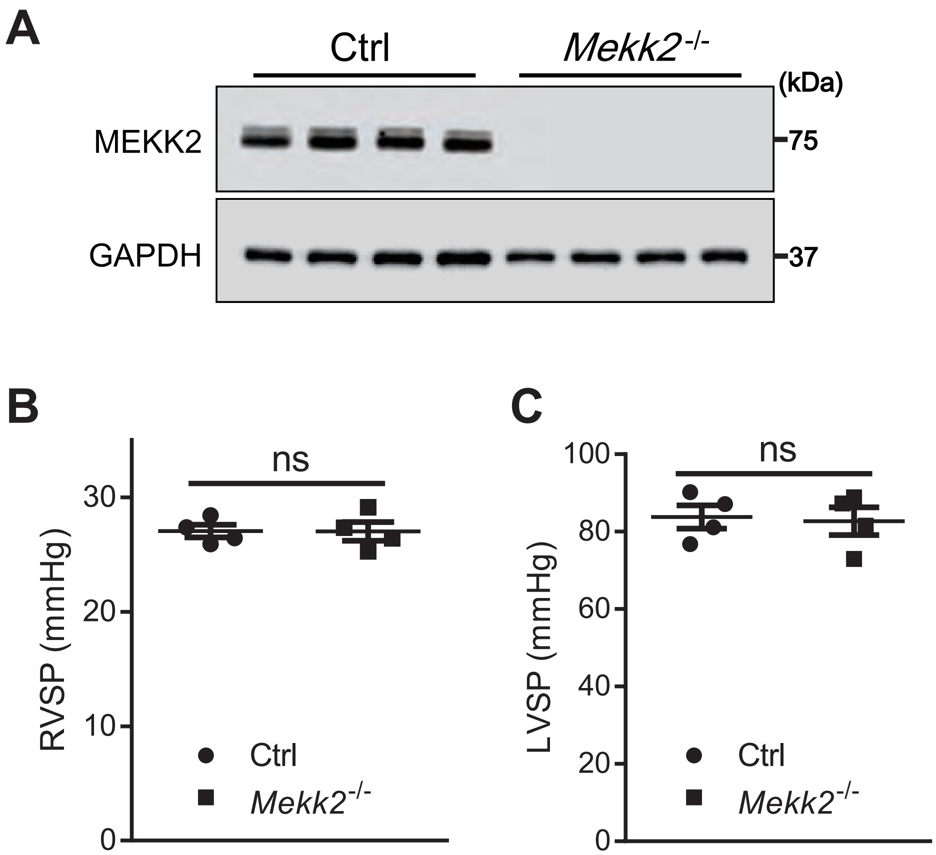

A
Ctrl
Mekk2-/-
(kDa)
MEKK2
75
GAPDH
37
B
ns
RVSP (mmHg)
30
20
10
0
Ctrl
Mekk2-/-
C
ns
LVSP (mmHg)
100
80
60
40
20
0
Ctrl
Mekk2-/-

**Figure S4**

**A**

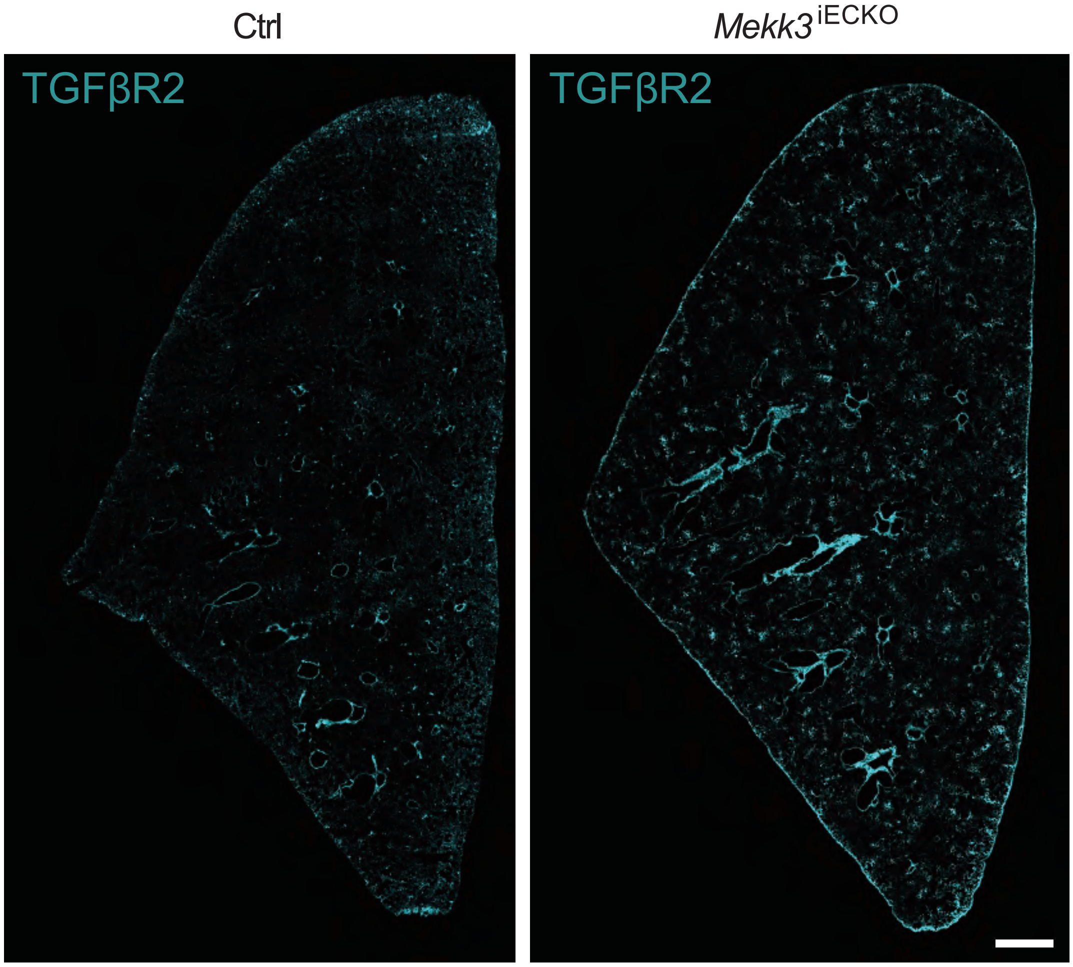

### Figure S5

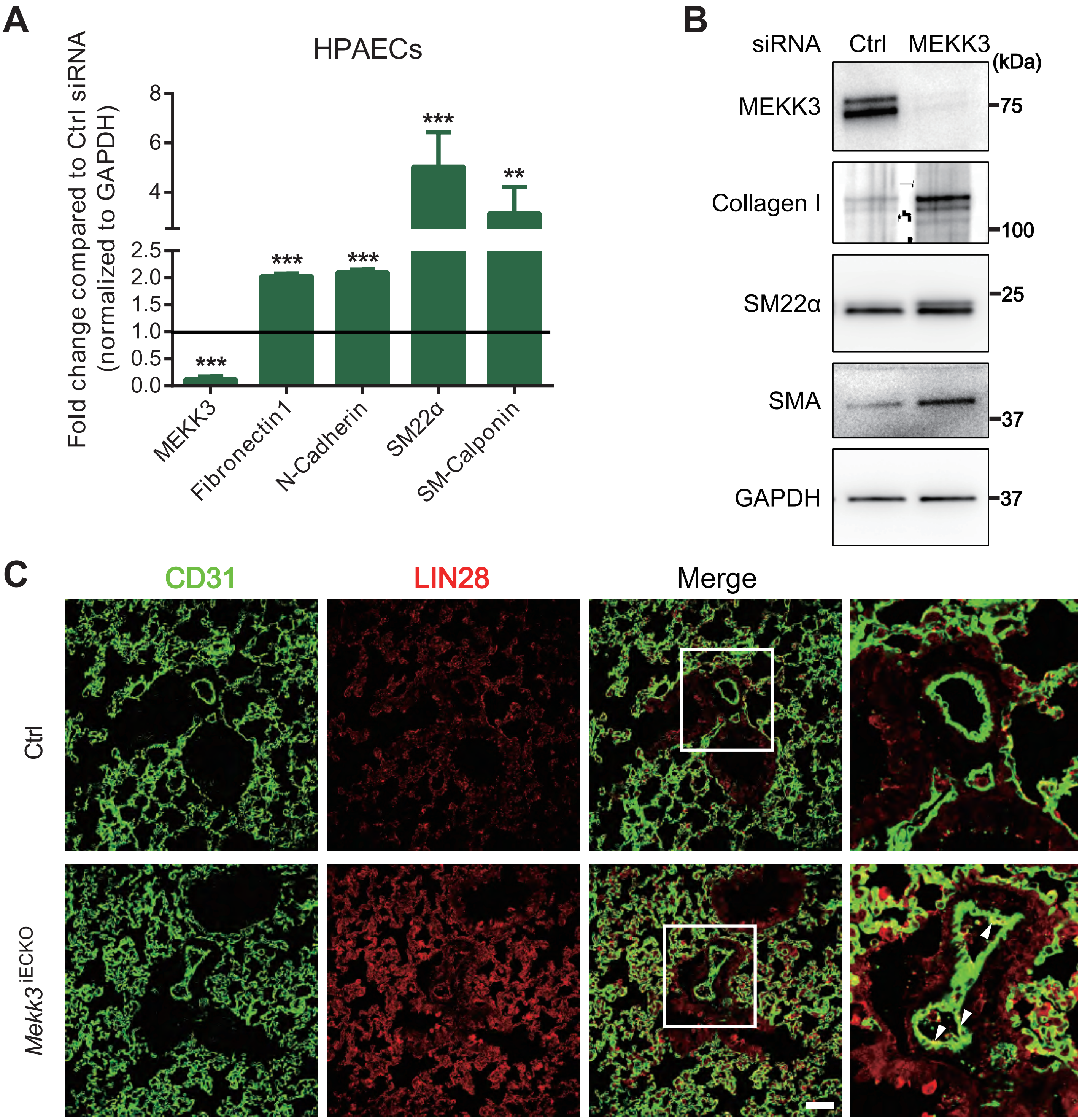

A
HPAECs
Fold change compared to Ctrl siRNA (normalized to GAPDH)
8
6
4
2.5
2.0
1.5
1.0
0.5
0.0
***
***
***
***
**
MEKK3
Fibronectin1
N-Cadherin
SM22α
SM-Calponin
B
siRNA
Ctrl
MEKK3
(kDa)
MEKK3
75
Collagen I
100
SM22α
25
SMA
37
GAPDH
37
C
CD31
LIN28
Merge
Ctrl
Mekk3 iECKO

### Figure S6

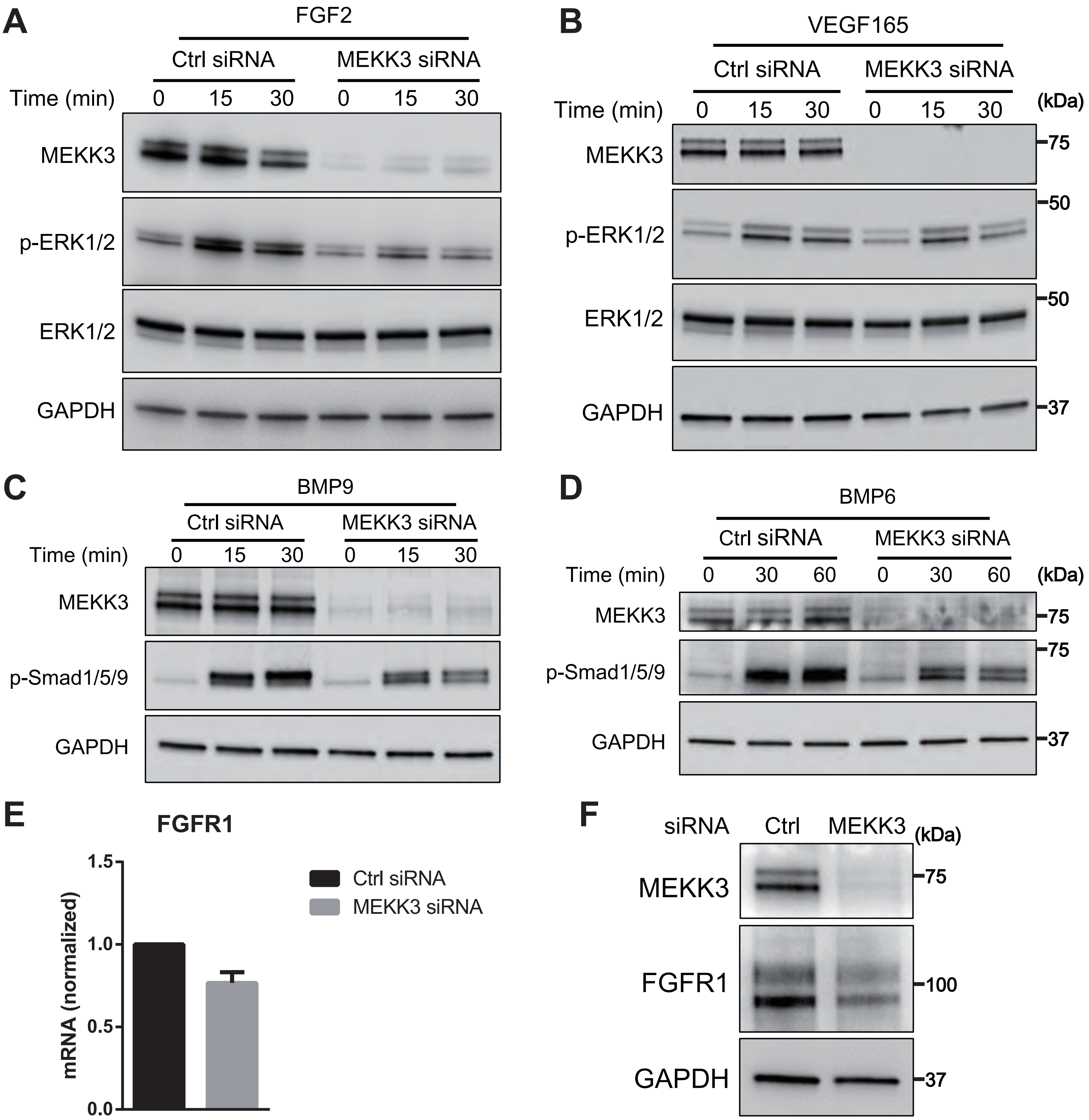
A
FGF2
Ctrl siRNA
MEKK3 siRNA
Time (min)
0
15
30
0
15
30
MEKK3
p-ERK1/2
ERK1/2
GAPDH
B
VEGF165
Ctrl siRNA
MEKK3 siRNA
Time (min)
0
15
30
0
15
30
(kDa)
MEKK3
p-ERK1/2
ERK1/2
GAPDH
75
50
50
37
C
BMP9
Ctrl siRNA
MEKK3 siRNA
Time (min)
0
15
30
0
15
30
MEKK3
p-Smad1/5/9
GAPDH
D
BMP6
Ctrl siRNA
MEKK3 siRNA
Time (min)
0
30
60
0
30
60
(kDa)
MEKK3
p-Smad1/5/9
GAPDH
75
75
37
E
FGFR1
mRNA (normalized)
1.5
1.0
0.5
0.0
Ctrl siRNA
MEKK3 siRNA
F
siRNA
Ctrl
MEKK3
(kDa)
MEKK3
FGFR1
GAPDH
75
100
37

### Figure S7

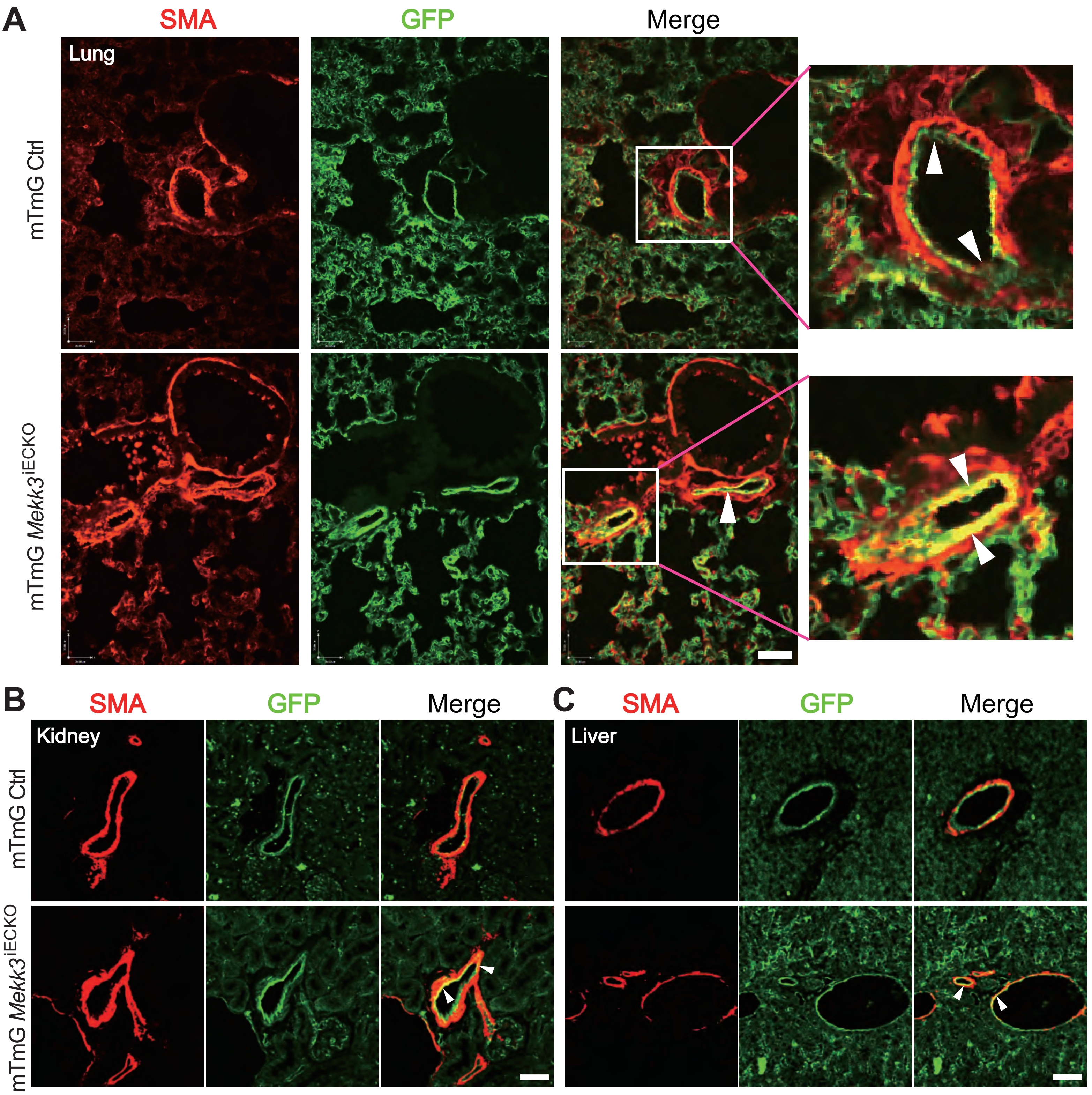
A
SMA
GFP
Merge
Lung
mTmG Ctrl
mTmG *Mekk3*$^{iECKO}$
B
SMA
GFP
Merge
Kidney
mTmG Ctrl
mTmG *Mekk3*$^{iECKO}$
C
SMA
GFP
Merge
Liver

#### Figure S8

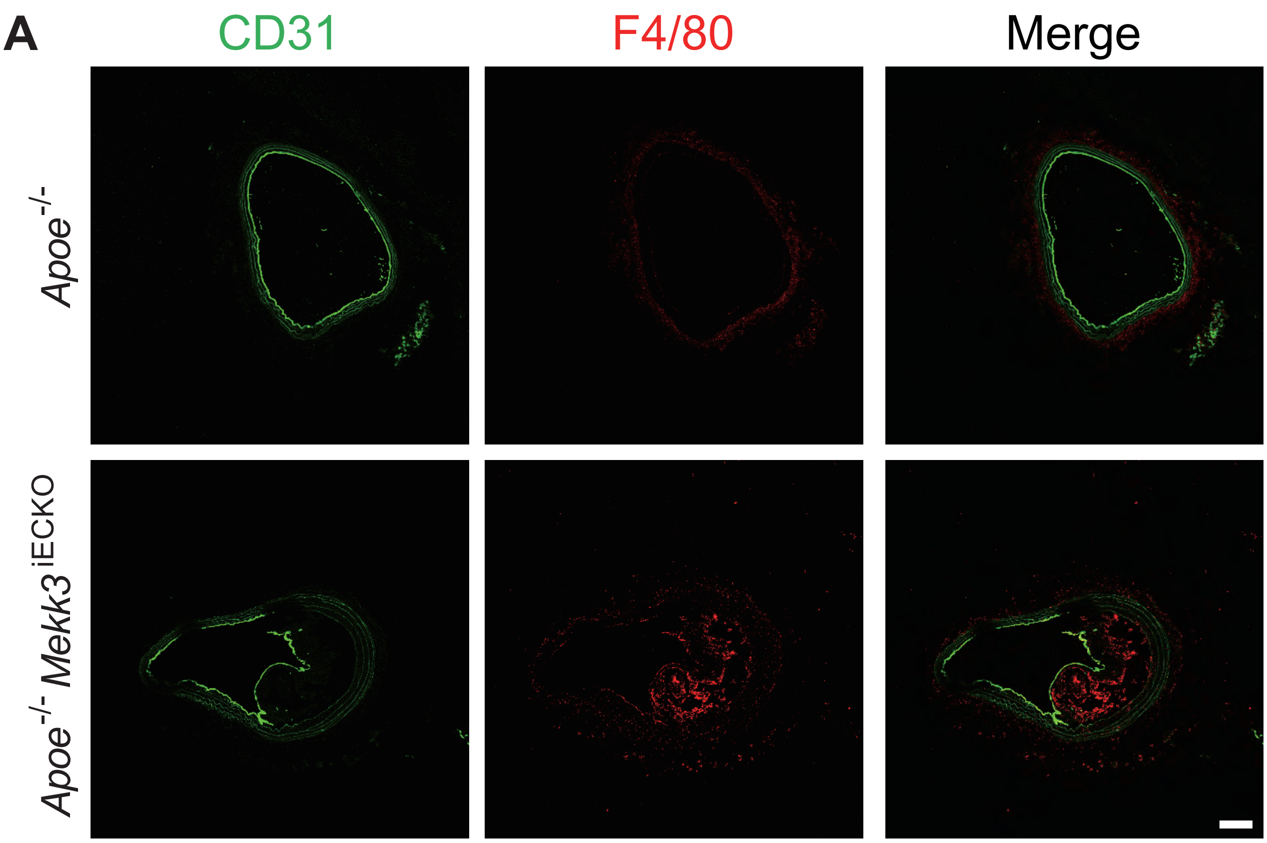

### Figure S9

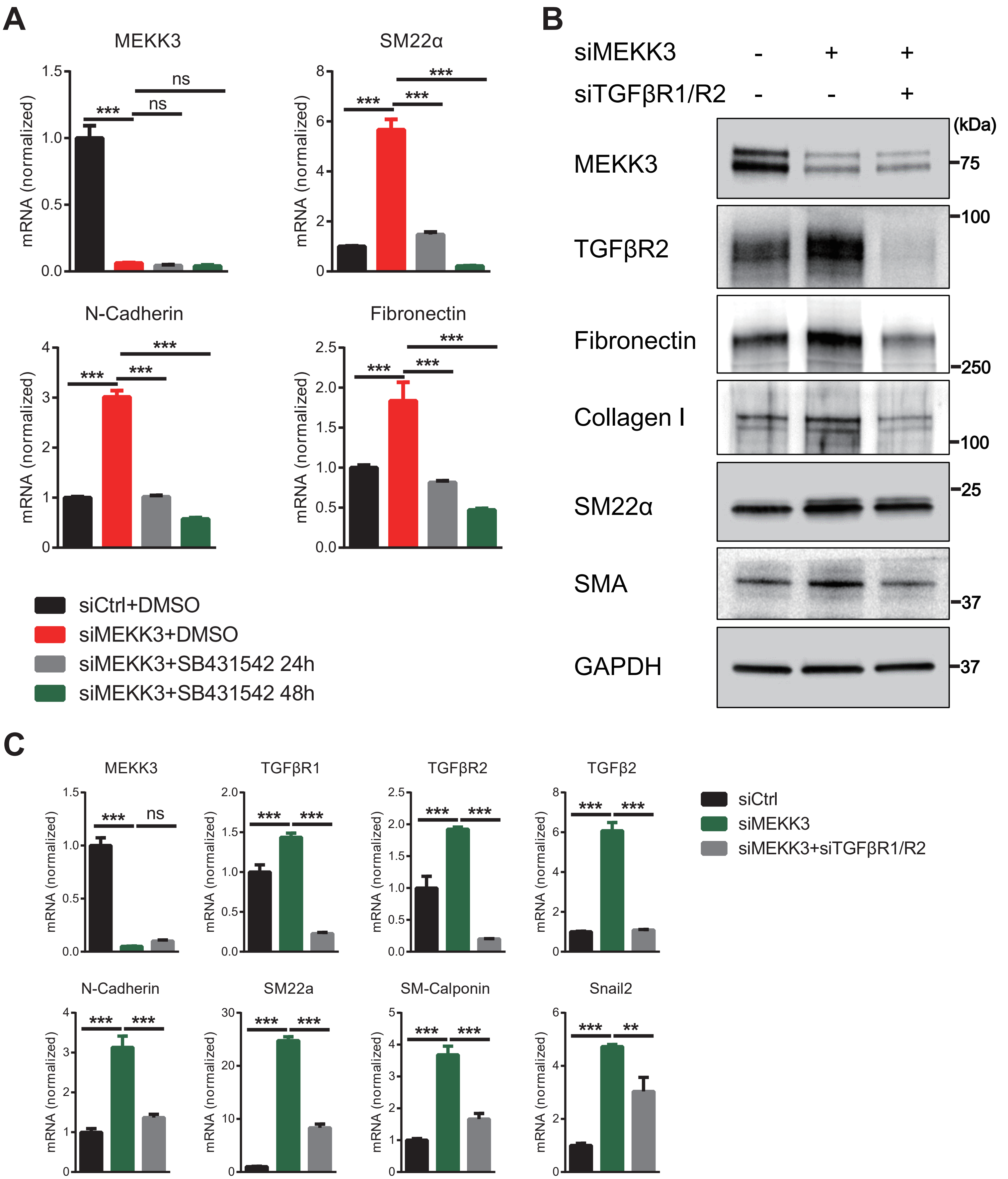
A
MEKK3
SM22α
N-Cadherin
Fibronectin
mRNA (normalized)
***
ns
siCtrl+DMSO
siMEKK3+DMSO
siMEKK3+SB431542 24h
siMEKK3+SB431542 48h
B
siMEKK3 - + +
siTGFβR1/R2 - - +
(kDa)
MEKK3
TGFβR2
Fibronectin
Collagen I
SM22α
SMA
GAPDH
75
100
250
100
25
37
37
C
MEKK3
TGFβR1
TGFβR2
TGFβ2
N-Cadherin
SM22a
SM-Calponin
Snail2
**
siCtrl
siMEKK3
siMEKK3+siTGFβR1/R2

**Supplemental Table 1. List of qPCR primers used.**

| Genes | Sequences (5' to 3' ) |
|---|---|
| hMEKK3-F | CAGACAGGAATACTCAGATCGGG |
| hMEKK3-R | TCTTCTGCCATCACTGTAGTCC |
| hGAPDH-F | GGAGCGAGATCCCTCCAAAAT |
| hGAPDH-R | GGCTGTTGTCATACTTCTCATGG |
| hFN1-F | GAGAATAAGCTGTACCATCGCAA |
| hFN1-R | CGACCACATAGGAAGTCCCAG |
| hCdh2-F | AGCCAACCTTAACTGAGGAGT |
| hCdh2-R | GGCAAGTTGATTGGAGGGATG |
| heNOS-F | TGATGGCGAAGCGAGTGAAG |
| heNOS-R | ACTCATCCATACACAGGACCC |
| hSnail2-F | TGTGACAAGGAATATGTGAGCC |
| hSnail2-R | TGAGCCCTCAGATTTGACCTG |
| hTGFβ1-F | GTACCTGAACCCGTGTTGCT |
| hTGFβ1-R | GTATCGCCAGGAATTGTTGC |
| hTGFβ2-F | ATGCGGCCTATTGCTTTAGA |
| hTGFβ2-R | GTTGGCATTGTACCCTTTGG |
| hTGFβ3-F | GCCTCAGTCTTTGGGATCTG |
| hTGFβ3-R | GTGTGAGCTGGGAAGAGAGG |
| hTGFβR1-F | CAGCTCTGGTTGGTGTCAGA |
| hTGFβR1-R | ATGTGAAGATGGGCAAGACC |
| hTGFβR2-F | TGAGTTCAACCTGGGAAACC |
| hTGFβR2-R | GGTTGATGTTGTTGGCACAC |
| hSM α-actin-F | CAAAGCCGGCCTTACAGAG |
| hSM α-actin-R | AGCCCAGCCAAGCACTG |
| hSM22α-F | GATTTTGGACTGCACTTCGC |
| hSM22α-R | GTCCGAACCCAGACACAAGT |
| hSM-calponin-F | CTGGCTGCAGCTTATTGATG |
| hSM-calponin-R | CTGAGAGAGTGGATCGAGGG |
| hLIN28a-F | AGCGCAGATCAAAAGGAGACA |
| hLIN28a-R | CCTCTCGAAAGTAGGTTGGCT |
| hLIN28b-F | CATCTCCATGATAAACCGAGAGG |
| hLIN28b-R | GTTACCCGTATTGACTCAAGGC |
